# Predicting human and viral protein variants affecting COVID-19 susceptibility and repurposing therapeutics

**DOI:** 10.1101/2023.11.07.566012

**Authors:** Vaishali P. Waman, Paul Ashford, Su Datt Lam, Neeladri Sen, Mahnaz Abbasian, Laurel Woodridge, Yonathan Goldtzvik, Nicola Bordin, Jiaxin Wu, Ian Sillitoe, Christine A Orengo

**Affiliations:** Institute of Structural and Molecular Biology, University College London, London, WC1E 6BT, UK; Department of Applied Physics, Faculty of Science and Technology, Universiti Kebangsaan Malaysia, Bangi, Malaysia

**Keywords:** COVID-19, human genetic variation, SARS-CoV-2: human protein interaction, protein structure complex, Functional Family, CATH database, protein binding affinity prediction, immunity

## Abstract

The COVID-19 disease is an ongoing global health concern. Although vaccination provides some protection, people are still susceptible to re-infection. Ostensibly, certain populations or clinical groups may be more vulnerable. Factors causing these differences are unclear and whilst socioeconomic and cultural differences are likely to be important, human genetic factors could influence susceptibility. Experimental studies indicate SARS-CoV-2 uses innate immune suppression as a strategy to speed-up entry and replication into the host cell. Therefore, it is necessary to understand the impact of variants in immunity-associated human proteins on susceptibility to COVID-19.

In this work, we analysed missense coding variants in several SARS-CoV-2 proteins and its human protein interactors that could enhance binding affinity to SARS-CoV-2. We curated a dataset of 19 SARS-CoV-2: human protein 3D-complexes, from the experimentally determined structures in the Protein Data Bank and models built using AlphaFold2-multimer, and analysed impact of missense variants occurring in the protein-protein interface region. We analysed 468 missense variants from human proteins and 212 variants from SARS-CoV-2 proteins and computationally predicted their impacts on binding affinities to SARS-CoV-2 proteins, using 3D-complexes.

We predicted a total of 26 affinity-enhancing variants from 14 human proteins implicated in increased binding affinity to SARS-CoV-2. These include key-immunity associated genes (TOMM70, ISG15, IFIH1, IFIT2, RPS3, PALS1, NUP98, RAE1, AXL, ARF6, TRIMM, TRIM25) as well as important spike receptors (KREMEN1, AXL and ACE2). We report both common (e.g., Y13N in IFIH1) and rare variants in these proteins and discuss their likely structural and functional impact, using information on known and predicted functional sites. Potential mechanisms associated with immune suppression implicated by these variants are discussed.

Occurrence of certain predicted affinity-enhancing variants should be monitored as they could lead to increased susceptibility and reduced immune response to SARS-CoV-2 infection in individuals/populations carrying them. Our analyses aid in understanding the potential impact of genetic variation in immunity-associated proteins on COVID-19 susceptibility and help guide drug-repurposing strategies.

## Introduction

The COVID-19 pandemic has caused a major global health and socioeconomic burden since 2020. Many countries are still experiencing an intermittent rise in the number of infections due to emergence of new Variants of Concern (VOCs) of SARS-CoV-2 and their sub-variants [1]. Although vaccines are now available, re-infection is common [2]. Various factors including ethnicity, age and clinical conditions have been proposed to be associated with an increased risk of infection [3–10]. In addition, increasing reports indicate that human genetic variation is a contributing factor for increased susceptibility and disease severity [11–13].

Potential drug targets include human host proteins which interact with SARS-CoV-2 [14, 15]. In 2020, Krogan group identified a total of 332 human proteins that interact with SARS-CoV-2 proteins, using affinity purification followed by mass spectrometry (AP-MS) [15]. Subsequently, the presence of additional human proteins interactors of SARS-CoV-2 was also revealed by other studies based on techniques such as yeast two-hybrid assay, anti-tag coimmunoprecipitation, tandem affinity purification, pull down, structure-based studies (X-ray, NMR), etc. (https://www.ebi.ac.uk/intact/home) [14, 16–19] and are made available via dedicated protein-interaction resources such as IntAct [20] and BIOGRID [21]. These studies indicate that interactor proteins in humans participate in a wide range of biological processes/pathways, including innate and adaptive immune pathways, lipid metabolism, cell adhesion, mRNA processing, among others [14, 15, 19, 22].

Innate immune suppression is known as one of the key characteristics of infections by SARS-CoV-2 and its VOCs. SARS-CoV-2 VOCs (namely Alpha, Beta, Gamma, Delta, and Omicron) are reported to exhibit increased interferon resistance as compared to the wild -type suggesting evasion of innate immunity is a driving force for SARS-CoV-2 evolution [23, 24]. Furthermore, inborn variation in immunity-associated genes is reported to trigger susceptibility to COVID-19 [25, 26]. For example, rare variants in the Toll-like receptor 7 gene have been associated with increased severity and susceptibility of the COVID-19 [12]. Likewise, various studies including those by the GeNOMICC-ISARIC consortium suggests association of both common and rare variants with increased severity of COVID-19 [27–32].

Though spike-ACE2 binding is the key entry mechanism used by SARS-CoV-2 for cell entry and infection, recent experimental studies indicate that SARS-CoV-2 also uses innate immune suppression as a strategy to speed-up entry and replication in the host cell [33, 34]. Interactions such as SARS-CoV-2:ORF9b-human:TOMM70 and SARS-CoV-2:NSP1-human:NUP98 have been associated with innate immune evasion [14, 34, 35]. Further experimental studies have suggested involvement of other human protein interactors of SARS-CoV-2 associated with the immunity-associated pathways (e.g., IFIH1, ISG15, IFIT2) [33, 34, 36].

In the case of SARS-CoV-2, spike-ACE2 is the most studied protein complex, where the impact of emerging variants in spike as well as natural human population variants in hACE2 have been computationally predicted by the Barton group and others [37–40]. Some of these predicted variants in ACE2 and spike protein, have also been validated experimentally [37, 40]. In this study, we focus on the impact of variants in immune-associated proteins and novel spike receptors (such as AXL and Kremen1) in humans on their binding to SARS-CoV-2 proteins. Such analyses could explain novel mechanisms by which SARS-CoV-2 proteins interfere with natural immune pathways and disrupt the system in humans.

Computationally predicted (docking-based) complexes for a subset of interactions identified from the Krogan study, are made available via several resources [41, 42]. The Beltrao group has designed a resource called mutfunc, which has predicted impacts of all possible single-amino acid substitutions in SARS-CoV-2 proteins, using known 3D structure data (http://sars.mutfunc.com/home, [43]). Likewise, Ascher group developed the COVID-3D resource, to analyse > 11,000 SARS-CoV-2 variants and predicted the impact of missense variants using spike-ACE2 complex [39]. More recently, a powerful AlphaFold2-based protein structure prediction method has been developed for modelling protein-protein complexes which facilitates modelling of interactions between SARS-CoV-2 and human proteins which have yet to be experimentally characterized, and with improved accuracy compared to other approaches [44–48].

In this study, we analysed the impact of missense coding variants in human and viral proteins occurring at protein-protein interface, using a curated dataset of 19 immune-associated SARS-CoV-2: human protein 3D-complexes, obtained from the Protein Data Bank [49] and models built using AlphaFold2-multimer [44, 46]. For the human proteins, we obtained population variants from various databases including gnomAD ([50] https://gnomad.broadinstitute.org) and GenomeAsia100k [51]. For the viral proteins, we obtained mutation data from ViralZone (https://viralzone.expasy.org/), CoV-Glue (https://cov-glue.cvr.gla.ac.uk/) and COG-UK (https://sars2.cvr.gla.ac.uk/cog-uk/) [52–54]. The impact of variants on binding affinity of the complexes was computationally predicted using a state-of-the-art program (mCSM-PPI2) [55]. The structural and functional impact of the predicted affinity-enhancing variants was analysed in the context of proximity to known functional sites such as protein-protein interface, ligand or substrate -binding sites and predicted sites identified using conserved positions in Functional Families in the CATH database (i.e., CATH-FunFams) [56–58]. CATH-FunFams represent functionally coherent groups i.e., members of a CATH-FunFam have high structural similarity and function [56]. Finally, the human proteins implicated in enhanced SARS-CoV-2 binding are mapped onto protein networks to understand biological pathways/processes associated with the network modules. We then studied whether these proteins are associated with CATH-FunFams that are enriched in small molecules or drugs from ChEMBL [59, 60].

In summary, we analysed the impact of missense coding variants occurring at protein-protein interfaces in a total of nineteen 3D complexes of human proteins and SARS-CoV-2 interactors. A total of 26 affinity-enhancing variants from 14 human proteins (namely TOMM70, IFIH1, IFIT2, ISG15, RPS3, PALS1, NUP98, RAE1, AXL, ARF6, KREMEN1, TRIMM, TRIM25 and ACE2) were predicted to enhance binding affinity to their interacting proteins in SARS-CoV-2. Our study sheds light on affinity-enhancing variants in immunity-associated proteins; their frequencies in gnomAD populations; their impact on protein structure and function and the populations more likely to be susceptible to COVID-19 infection. We provide computational evidence that the predicted affinity-enhancing variants in human proteins could promote binding to SARS-CoV-2 proteins, instead of their natural protein partners or substrates in immune pathways, thereby hampering the normal antiviral activity and leading to increased susceptibility. Protein Functional families associated with three proteins (IFIH1, AXL and ARF6) are associated with small molecule inhibitors and their potential applications in drug-repurposing in discussed.

## Materials and methods

### Compilation of interactors associated with SARS-CoV-2 immunity

The dataset of human proteins interacting with SARS-CoV-2 proteins, was compiled using the COVID-19 UniProtKB resource ([61]; https://covid-19.uniprot.org/uniprotkb?query=*) and IntAct database ([20]; https://www.ebi.ac.uk/intact/home). IntAct provides a COVID-19 dataset of SARS-CoV-2: human protein interactors which is based on interactions reported from experimental studies. For every protein-protein interaction, IntAct assigns an MIscore (ranges from 0 to 1) based on (i) the type of the experimental detection method (ii), the number of associated publications and (iii) the interaction types (such as direct association, physical association). The IntAct recommended MIscore threshold of 0.45 was used to exclude low-confidence interactions [62].

Thus, the dataset of a total of 536 high-confidence (MI-score > 0.45) interactions are considered for subsequent analyses. We further filtered immunity-associated human proteins by mapping specific GO terms associated with immunity using UniProt (i.e.,GO:0002250 GO:0002218, GO:0002376, GO:0045087, GO:0045089, GO:0060337, GO:0050776 and GO:0006955)[61, 63]; and by mapping the UniProt IDs to InnateDB database (http://innatedb.sahmri.com/ [64]. We also curated available literature-based evidence specifying COVID-19-associated immunological role associated with interacting pair of proteins in dataset.

### Functional families in the CATH database and conserved functional sites

The CATH database provides a hierarchical structural classification of protein domains into Class (C), Architecture (A), Topology (T) and Homologous Superfamily (H). In CATH, protein domains are classified into superfamilies where there is strong evidence of an evolutionary relationship via structure and sequence similarity [65, 66]. Within each superfamily, sequences are sub-classified using an entropy-based method to segregate functionally coherent subgroups known as Functional Families (CATH-FunFams) [56]. The conserved sites obtained from CATH-FunFams have been shown to be enriched in known protein functional sites [56, 57].

For the dataset of shortlisted human proteins and their SARS-CoV-2 interactors, we identified CATH-FunFams, and subsequently conserved sites as follows:

- *Identification of CATH-FunFams:* We scanned the sequences of human and their interacting SARS-CoV-2 proteins against HMMs of FunFams in CATH v4.3 (https://www.cathdb.info/), using HMMsearch (e-value 1e^-3^) [67]. We then processed the output of HMMsearch using cath-resolve-hits, an in-house tool built to obtain the best non-overlapping set of domain matches [67, 68].
- *Identification of conserved sites using CATH-FunFams*: Using the matching CATH-FunFams, we identified conserved sites using Scorecons, an entropy-based method [69]. The multiple sequence alignment (MSA) program, namely MAFFT is used to construct an MSA from seed sequences within a FunFam [68]. The Scorecons program is then applied to each MSA to determine an overall measure of sequence diversity called Diversity of Positions score (DOPs). DOPs captures the amount of diversity in an MSA by considering all the different conservation scores, and their frequencies, and provides a value from 0 (i.e., zero diversity) and 100 (i.e., high diversity). Only MSAs with a DOPs score ≥ 70 were considered for further analyses. The Scorecons program also provides the degree of conservation of each position in the MSA. Thus, for each column in a CATH-FunFam based MSA, the Scorecons program provides a conservation score ranging from 0 (i.e., not conserved) to 1 (i.e., completely conserved). The sites belonging to alignment positions with Scorecons-based score >= 90 are used for analyses and are referred to as Scorecons90 in this manuscript.

### Compilation of missense coding variants in human and SARS-CoV-2 proteins

- *Human protein variants:* For the human genes, missense coding variants from canonical transcripts were obtained from the Genome Aggregation Database (gnomAD, v2.1.1; the recommended version for coding region analyses) (https://gnomad.broadinstitute.org/) [50]. We compiled ancestry (i.e., ethnic population) information available from gnomAD, using VarSite [70]. GnomAD provides ancestry for the following populations: African/African American (afr), American Admixed/Latino (amr), Amish (ami), Ashkenazi Jewish (asj), East Asian (eas), South Asian (sas), Finnish (fin) and Non-Finnish European (nfe). If individuals did not unambiguously cluster with any of these populations in a principal component analysis (PCA), gnomAD classifies them as “other” (oth). GnomAD v2 also provides sub-continental information for the East Asian cohort (Japanese, Koreans) and Non-Finnish European (Bulgarian, Estonian, Swedish, North-Western European, Southern European) populations.
- *SARS-CoV-2 protein variants:* For each of the associated interactor proteins in SARS-CoV-2, a non-redundant set of mutations in strains of SARS-CoV-2 (including VOCs and Variants of Interest) was compiled from resources such as ViralZone [54], COG-UK ([52] https://sars2.cvr.gla.ac.uk/cog-uk/), and CoV-Glue (https://cov-glue-viz.cvr.gla.ac.uk/) [53].

### Functional site data from CATH-FunVar (Functional Variation) protocol

All gnomAD missense variants in the interactor human genes were processed using a modified version of the in-house FunVar protocol to identify variants occurring near known and predicted functional sites. Predicted sites comprise CATH-FunFam-based conserved residues, i.e., highly conserved residues identified by the Scorecons program (i.e., Scorecons90 sites, as described above).

Known functional sites were compiled from existing resources and include: known ligand and nucleic acid binding sites from BioLip [71]; protein-protein interface (PPI) residues from PDBSum [72]; catalytic sites annotated in M-CSA and VarMap [73, 74] and annotated functional sites in UniProt resource.

Spatial proximity of each gnomAD variant to these sites was found by mapping each variant to a representative domain from the corresponding FunFam in CATH v4.3. Functional sites were similarly mapped to FunFam representative domains, allowing detection of variants occurring on or near (within 5Å) of any of the functional sites.

For each variant, the Grantham score (https://gist.github.com/danielecook/501f03650bca6a3db31ff3af2d413d2a) was calculated to identify variants having a significant change in physico-chemical properties (such as volume, polarity), as compared to that of wild-type residues.

Finally, each variant was assigned a simple functional impact score (from 1 to 5) by counting each of the impacts-scoring 1 for each of: high Grantham score; variant is a catalytic site; variant lies on a known functional site; variant near (5Å) a known site; variant is on a conserved predicted (Scorecons90) site. Additionally, CATH-FunVar reports impact scores from CADD [75] and SIFT [76] .

### Three-dimensional (3D) structures of complexes

3D structures of complexes are available for 10 interactions as follows-human:TOMM70-SARS-CoV-2:ORF9b [PDB ID: 7KDT], human:ISG15-SARS-CoV-2:PLpro [7RBS], human:RPS2-SARS-CoV-2:NSP1 [6ZMT], hRPS3-SARS-CoV-2:NSP1 [6ZMT], human:Ubiquitin-SARS-CoV-2:Plpro [7RBR], human:APOA1-SARS-CoV-2:ORF3a, human:PALS1-SARS-CoV-2:E, human:NUP98-SARS-CoV-2:ORF6 [7VPH] and humanRAE1-SARS-CoV-2:ORF6 [7VPH] and human:ACE2-SARS-CoV-2:spike [wild-type (6M0J, 7A95); Alpha (7EDJ), Beta (7V7Z), Gamma (7V83), Delta (7V89), Omicron (7T9K), BA.1 (7XO6), BA.2(7XB0, 7XO8), BA.3(7XB1)].

For the remaining interactions with no available structures of complexes in the PDB, we predicted models using AlphaFold2-ptm and AlphaFold2-multimer(v1) [44, 46], as described below.

- *Modelling complexes using AlphaFold2-ptm and AlphaFold2-multimer:* Prior to modelling protein-protein complexes, we excluded some interactions from the modelling procedure due to the following reasons – we excluded proteins of very short lengths such as ORF3b (22aa residues long) and proteins for which high-quality models are not built by AlphaFold2 (https://alphafold.ebi.ac.uk/). We modelled the remaining complexes using the AlphaFold2-ptm and alphafold-multimer(v1) protocols, which were made available in March, 2022 ([44, 46, 77]; https://github.com/sokrypton/ColabFold]. We built models using both the AlphaFold2_ptm and AlphaFold2-multimer(v1) methods and then selected a model from one of these methods, whichever had the best interface quality (see results, table 1). High-confidence models were chosen where models have overall pLDDT (predicted local difference distance test) ≥ 70 as well as pTM-Score (predicted TM-score) ≥ 70 [44, 46]. We further filtered complexes on the basis of the interface quality i.e., interface-pLDDT (i.e., those with <70 were excluded) and interface-PAE (predicted alignment error >10 were excluded) and by manually inspecting the domain interface regions (i.e., excluded models where we observed erroneous overlapping/entangled interface). We performed additional quality checks such as verifying interface stability score using PIZSA method (which calculates protein interaction Z-Score; > 1.5 indicates stable interface) [78] and predicted binding affinity of the complexes by the PRODIGY method [79]. The resultant high-confidence models predicted in this study along with their quality metrics are given in results section, Table 1.

**Table 1:**
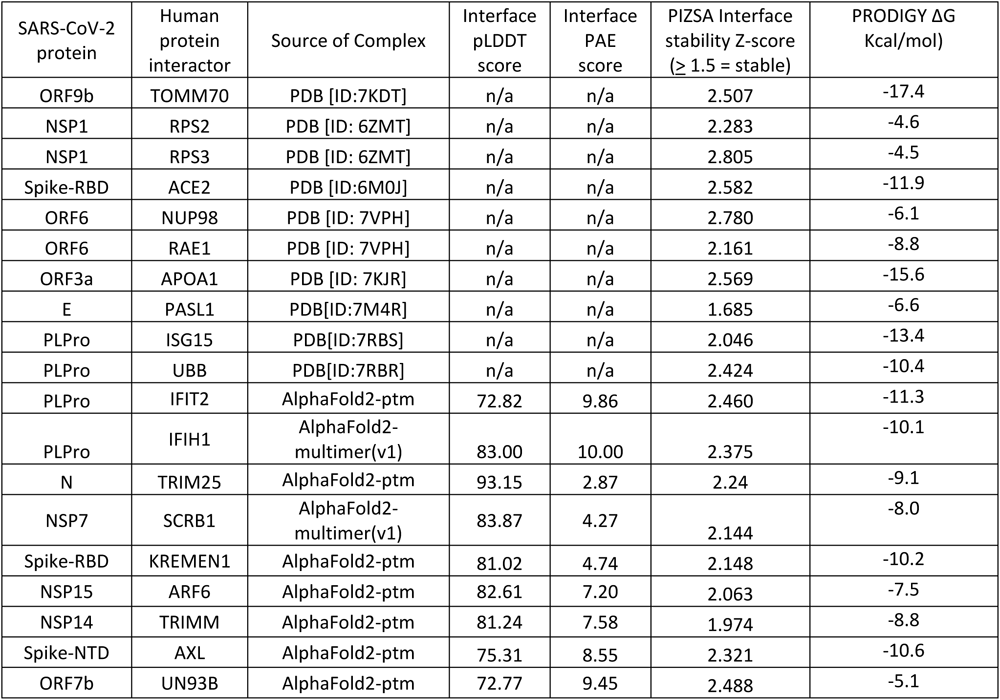
Summary of human: SARS-CoV-2 protein complexes used for the study. The table lists 3D complexes used in this study. We outline the source, quality metrics such as Interface pLDDT (≥70), Interface PAE (≤10), PIZSA Interface stability Z-score (≥1.5 indicates stable interface) and PRODIGY binding energy (ΔG ≤ -4.3 Kcal/Mol, cutoff chosen based on experimental binding energy values reported in PRODIGY reference. The literature supporting biochemical evidence of protein/domain interaction is also cited in Supplementary file 2. For modelled complexes, we first built models using both the AlphaFold2 (ptm method) or AlphaFold2-multimer(v1) and then selected model from one of these methods, whichever had the best interface quality.

- Extraction of interface residues using complexes After the selection of the complexes, we extracted interface residues as follows:
- Directly-contacting (DC) residues For the PDB complexes, we extracted directly-contacting (DC) interface residues made available by PDBSum [72]. PDBSum calculates the protein-protein interfaces using the NACCESS program (http://www.bioinf.manchester.ac.uk/naccess/). For the modelled complexes, we extracted interface residues by selecting residues from interacting chains with heavy atom distances <=4Å) [80, 81]. Human protein missense variants in gnomAD that occur in DC interface residues are referred to as ‘DC-variant’ residues.
- Secondary shell residues Residues that occur within 5Å from the DC interface are considered to be residues in the secondary shell and variants at these residue positions are likely to influence binding [38, 82] and are referred to as ‘DCSS-variants’.

### Predicting the impact of variants in human and viral proteins on the binding affinity of the complexes

For the filtered dataset of complexes (experimental and predicted), we applied the mCSM-PPI2 program [55] to identify human and viral missense variants which could significantly impact binding-affinity of interacting proteins. We analysed the impact of missense variants reported in gnomAD that occur at directly contacting interface (DC-variant) and secondary shell (DCSS-variant) residues.

The mCSM-PPI2 program, developed by the Ascher lab, was shown to be the top-performing method when compared to 26 other methods in CAPRI (round 26) blind tests [55]. mCSM-PPI2 is based on a graph-based structural signature framework with evolutionary information, inter-residue non-covalent interaction networks analysis plus computed energetic terms, providing an optimised overall prediction performance. It was used to predict change in binding affinity (i.e., mCSM-PPI2 ΔΔG^Affinity^ in kcal/mol) for each mutation.

A positive mCSM-PPI2 ΔΔG^Affinity^ score (> 0 Kcal/mol) indicates that the mutation is stabilising the interaction whereas a negative ΔΔG score (< 0 Kcal/mol) indicates a destabilising effect [55]. Where available, we compiled evidence from experimental studies reporting experimental mutagenesis and binding affinity kinetic assays [14, 37] to choose our ΔΔG thresholds. For example, in the case of spike-ACE2 complex, the mutations K26R in ACE2 and S477N in the spike protein are reported to increase the binding affinity of the spike-ACE2 complex using kinetic assays [37, 38], and these are predicted to have mCSM-PPI2 ΔΔG affinity of 0.12 Kcal/mol and 0.5 Kcal/mol respectively. Likewise, experimental alanine scanning mutagenesis studies in the ORF9b-TOMM70 complex indicates that a E477A mutation in TOMM70 and a S53A mutation in ORF9b significantly reduced the binding affinity of the interaction [14], and are predicted to have a mCSM-PPI2 ΔΔG^Affinity^ of < -0.5 Kcal/mol. Therefore, we used the confidence cut-offs of ΔΔG^Affinity^ ≤ -0.5 Kcal/mol (for destabilising; affinity-reducing) and ≥ 0.5 Kcal/mol (for stabilising; affinity-enhancing), to analyse the structural impact of gnomAD variants in this study, while we also provide a catalogue of other mutations predicted to have scores ranging from 0 to 0.490 Kcal/mol (see results section). Through *in silico* saturation mutagenesis, we further confirmed that the mCSM-PPI2 method does not show bias towards predicting positive ΔΔG^Affinity^ scores and corresponds well to observed changes in amino acid properties for mutated residues.

*Analysis of predicted affinity-enhancing variants*: As the affinity-enhancing variants could be associated with increased risk of susceptibility/infection, we closely examined their 3D structural impact on molecular interactions and characterised these variants in the context of known and predicted functional sites (See FunVar section above) and using the UniProt’s site annotations. Variants are mapped on 3D-complexes and visualized using UCSF Chimera [80]. We further analysed their associated population allele frequencies in gnomAD as well as other databases such as Indigenomes [83], SweGen [84], GenomeAsia100k [51], jMORP [85] and the NIH-funded research hub called All of Us ([86] https://databrowser.researchallofus.org/). Finally, we analysed pathogenicity scores by mutpred2, CADD and SIFT for affinity-enhancing variants. These methods were chosen as the study by Pejaver group showed that Mutpred2 performs better than other predictors while CADD and SIFT were the second and third best performing tools [87].

### Network mapping and enrichment analysis

- **Identification of network modules** The human proteins containing affinity-enhancing variants with putative impact on SARS-CoV-2 binding, were mapped to ConsensusPathDB (CPDB) and STRING database (STRINGdb) protein-protein interaction networks [88, 89]. Interactions from STRINGdb were filtered to include only those with confidence values >=0.2 to optimize signal-to-noise ratio. A module detection algorithm (M1) was applied to the network using the MOdularising NEtwork Toolbox that adopts a multiresolution approach to combine optimization algorithms to improve modularity [90].

- Enrichment analyses Pathway enrichment analysis was performed using g:profiler (https://biit.cs.ut.ee/gprofiler/gost, [91]), a public webserver using Ensembl and Ensembl Genomes to identify significantly enriched terms in Gene Ontology (GO) biological processes, KEGG and Reactome databases [92–94]. Pathways were ranked by significance (p<0.01) and the most significant term per database for the protein or each module was taken as the representative pathway. Where necessary, Ensembl identifiers with the most GO annotations were selected according to the default g:profiler function. Values and pathways for STRINGdb and CPDB modules were retained.

### Identification of druggable functional families in CATH database

The human proteins with affinity-enhancing variants, were mapped to the CATH functional families (CATH-FunFams) that are linked with small molecule information from ChEMBL [59, 60]. For the associated human proteins, the druggability score of the protein-protein interface region was analysed using the CavityPlus program [95].

The methodology used in the study is summarised in Figure 1.

**Figure 1:**
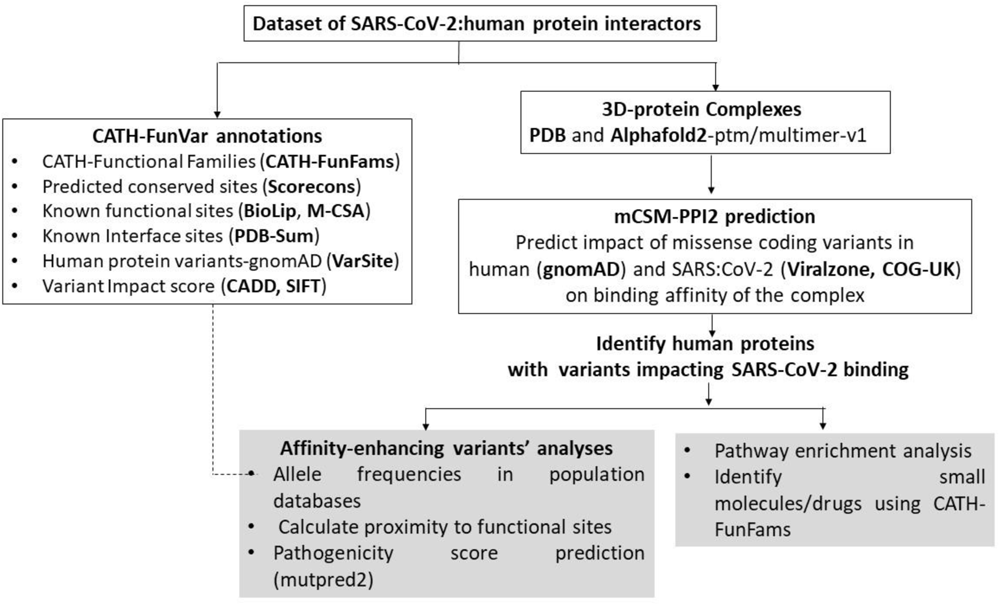
Flow-chart of the methodology used in this study. The dataset of SARS-CoV-2: human protein interactors is analysed using variants from gnomAD (in case of human proteins). In-house developed CATH-FunVar (Functional Variation) pipeline was used to extract all annotations on known and predicted functional sites. For the missense coding variants from gnomAD, the impact of binding affinity of the complex was analysed using mCSM-ppi2 program. Affinity-enhancing variants analysed using functional sites, and population data from gnomAD. The human proteins predicted to contain affinity-enhancing variants are used for pathway enrichment and identification of CATH functional families (CATH-FunFams) linked with small molecules/drugs from ChEMBL.

## Results

### Dataset of SARS-CoV-2: human protein interactors, 3D-complexes and missense variants

As previously described (see Methods), from 536 high-confidence (MIscore ≥ 0.45) protein-protein interactions, we curated 94 human proteins involved in SARS-CoV-2 infection and immunity by identifying those with immunity-associated GO terms. For the curated dataset of 94 human proteins, we compiled information from multiples sources: the interactor protein in SARS-CoV-2, IntAct MIscore [20], UniProt accession IDs [61], immune-associated GO terms, literature evidence linked to the interaction in IntAct, literature-based evidence for COVID-19 immune association, CRISPR-association (obtained from BIOGRID-ORCS; https://orcs.thebiogrid.org/ [21]) and gene-expression (obtained from SARS-COVIDB; https://sarscovidb.org/ [96]). This information is provided in Supplementary files 1 and 2.

In total, the 94 human proteins were associated with 110 SARS-CoV-2:human interactions (Supplementary file 2). Experimental 3D structures for 10 interactions were available in the PDB, with a further 9 high-quality models predicted using AlphaFold2-multimer/ptm method, as summarised in Table 1. Thus, a total of 19 protein 3D structural complexes were used for subsequent analyses.

For the human proteins from this curated dataset of 19 complexes (Table 1), we analysed a total of 468 missense variants from gnomAD; that occur at directly contacting (DC) residues in the interface of the complexes (DC-variants, 131 in total) as well as those that occur within 5Å from the DC residues i.e., secondary shell (DCSS-variants, 337 in total).

A total of 26 variants from 14 human proteins, are predicted to significantly enhance binding affinity to their SARS-CoV-2 protein partners by the mCSM-ppi2 program (ΔΔG^Affinity^ > 0.5 kcal/mol), as detailed in the Table 2.

**Table 2:** 14 Human proteins with variants predicted to have significant impact on binding affinity to SARS-CoV-2 interactor proteins (by mCSM-PPI2 program [55]). The first column lists the name of complexes in the following format-human protein (SARS-CoV-2 interactor protein). For every complex, the variants in human proteins that are predicted to enhance binding affinity with ΔΔG^Affinity^ score ≥0.5 Kcal/mol are indicated in bold. The variants that are predicted to increase binding affinity but with lower scores ranging from 0 < ΔΔG < 0.49 Kcal/mol are provided in the last column. * Indicates that the ACE2 variants reported in our study are also noted to enhance binding affinity in previous studies [reviewed in 45].

| Name of the complex:<br>human (SARS-CoV-2 protein) | Affinity-enhancing variants in human proteins<br>($\Delta\Delta G^{\text{Affinity}} \geq 0.5$ kcal/mol)<br>(Stabilizing mutations) | Affinity-reducing variants in human proteins<br>($\Delta\Delta G^{\text{Affinity}} < -0.5$ kcal/mol)<br>(de-stabilizing mutations) | Other affinity-enhancing variants in human proteins<br>($0 < \Delta\Delta G < 0.49$ Kcal/mol) |
| --- | --- | --- | --- |
| hTOMM70 (ORF9B) | **V556L, K576R, A591T, V514I, A483T** | I412T, Q477G, H515Q, L584M, I554T, D589N, L111P, F155L, G530S, D397N | A582V |
| hISG15 (PLPro) | **L121Q** | L60V, G128V, L28M, F122L, P130T, E127G, M23V, N151D, L85P, M23T, R155W, N151K, R155Q | S21N, Q55R, Q118R, V148L, A30V, D56Y, Q31H |
| hIFIH1 (PLpro) | **Y13N, S16L** | - | L120F, F107S, L121F, M95 |
| hIFIT2 (PLpro) | **L373F, K221E, A319S, A319T, L373F, Y383F** | N300K, R376Q, R406S, L375V, R376P, Q384K, L323P, G398V, F380L | Q381H, R292K, P190Q, A391V |
| hRPS3 (NSP1) | **V164I, I99F** | R106C, L86H, Y34H, V153A | E81Q, A52T, I223V, A114P, |
| hNUP98-hRAE1(ORF6) | **NUP98: T190S**<br>**RAE1: A275V** | - | NUP98: K185R, I162V.<br>RAE1: 275-CANCER, S311L |
| hARF6 (NSP15) | **L166F** | I29V, L33V, Y78C | N56H, D68H, I96F |
| hTRIM25 (N) | **A466T** | V470M, F615L | C475Y, P490L, I457V, A471S, K458E, C506S |
| hTRIMM (NSP14) | **I105F** | - | Q160R, H223Y, K146R, A144E, I13F |
| hAXL (spike-NTD) | **V38M** | L54I, Q57K | A79E |
| hKremen1 (Spike-RBD) | **Y66H, V189L** | Y98C, C200Y, C86Y, E187K, V93M, E187K, V93M, H196L, Y167C, Y107H, G166D, W108R | - |
| hACE2 (spike-RBD) | **G326E\*** | G352V, D355N, L585P | A501T*, K26R*, S19P*, K26E* |
| hPALS1(E) | **L321F** | - | - |

For the affinity-enhancing variants (ΔΔG^Affinity^ scores ≥ 0.5 Kcal/mol) from the 14 human proteins (Table 2), we analysed their impact on protein structure and function and their allele frequency distribution related to distinct populations in the gnomAD database and other databases including GenomeAsia100K (see methods). Most of the predicted affinity-enhancing variants are rare in gnomAD populations, while a few affinity-enhancing variants are observed to be common in genes such as IFIH1 and ISG15 (See figure in Supplementary file 3).

For the 14 human proteins, affinity-enhancing variants were analysed in the context of proximity to known and predicted functional sites (Supplementary file 4). The potential mechanisms associated with suppression of the immune system implicated by these variants are discussed in the following section, using some complexes to illustrate our approach.

### Structure-function analyses of predicted affinity-enhancing variants

#### 1. Impact of human TOMM70 variants on SARS-CoV-2: ORF9b binding

TOMM70 protein is one of the major human import receptors in the translocase of the outer membrane (TOM) complex. It recognizes and mediates the translocation of mitochondrial preproteins from the cytosol into the mitochondria in a chaperone(HSP90)-dependent manner [97]. It is involved in activation of the innate immune system [GO:0002218] and interacts with ORF9b, which is a key viral innate immune antagonist in SARS-CoV-2 [14, 97, 98]. A study by the Krogan group revealed that TOMM70 is a high-confidence interactor of SARS-CoV-2 ORF9b indicating that binding of ORF9b to the C-terminal domain of TOMM70 is associated with suppression of the innate immune response [33]. We analysed the impact of missense variants using the experimental structure of this complex (PDB ID: 7KDT [14]).

#### Affinity-enhancing variants: structural-function impact and population distribution

Three DC (directly contacting)-variants in TOMM70 (V556L, K576R, A591T) and two DCSS-variants (V514I, A483T) were predicted to significantly increase affinity (ΔΔG^Affinity^ > 0.5 Kcal/mol) (Table 3 and Figure 2).

**Figure 2:**
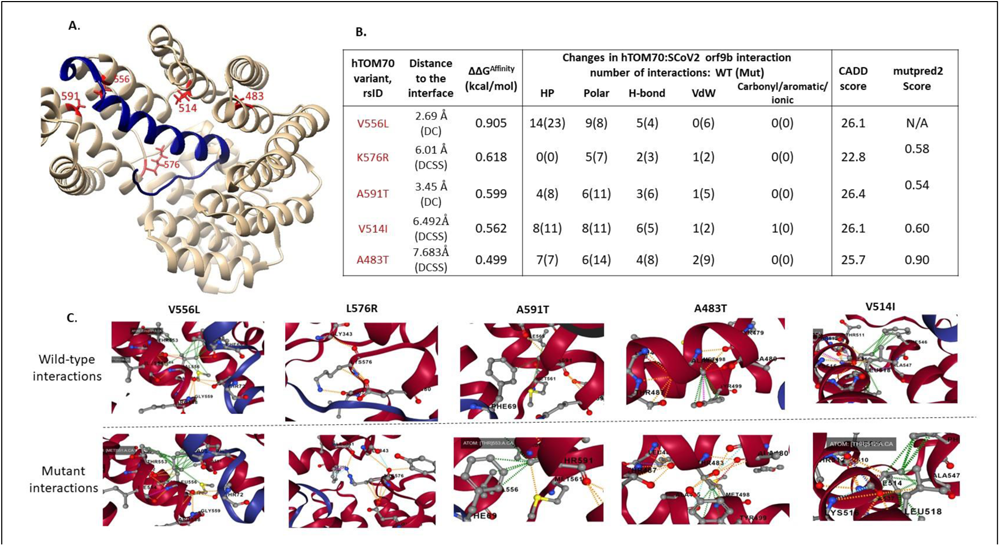
The structure of human TOMM70 in complex with SARS-CoV-2 ORF9b [PDB ID: 7KDT]. (A) ORF9b-TOMM70 complex: SARS-CoV-2 ORF9b (blue) interacts with the C-terminal domain of human TOMM70 (tan). ORF9b binds at the substrate-binding pocket in TOMM70. The structural locations of variants (with ΔΔG^Affinity^ ≥ 0.5 Kcal/mol) are shown in red. These include three DC-variants (V556L, K576R, A591T) and two DCSS-variants (V514I, A483T). (B) The structural impact of DC-variants on atomic interactions at the interface, are shown. HP: Hydrophobic (green), H-bond: hydrogen bond (red), VDW: Van-der-Waals (blue), Polar (orange) (C) Effects of affinity-enhancing variants in TOMM70 are shown in detail (figures in 2C source: mCSM-PPI2).

**Table 3:** Details of variants in TOMM70 that impact binding to ORF9b. For each variant, SNP ID (i.e. rsID), amino acid mutation, mCSM-PPI2 prediction, functional site annotations and population in gnomAD (in which these variants are present) are indicated. Functional sites are abbreviated as follows-DC_PPI: directly contacting interface site, DCSS_PPI, secondary shell residue from the DC interface site, Ligsite: indicates occurrence in a known ligand binding site and near_ligsite indicates sites that are proximal to a known ligand (HSP90) binding site. Variants with Mutpred2 pathogenicity score ≥ 0.50, are indicated.

| Variant ID | rsID | Missense variant in gnomAD | mcsM-ppi2-prediction (kcal/mol) | Functional site annotation | GnomAD populations [and their allele frequency -AF] |
| --- | --- | --- | --- | --- | --- |
| 3-100086895-C-T | rs1306989616 | p.Val556Leu | **0.905** | DC_PPI | African/AA [AF: 0.00007] |
| 3-100084508-T-C | rs1370508158 | p.Lys576Arg | **0.618** | DC_PPI<br>Mutpred2_0.58 | South Asian [0.00003] |
| 3-100084464-C-T | rs756063544 | p.Ala591Thr | **0.599** | DC_PPI<br>Mutpred2_0.53 | Latino/AA [0.00008]<br>AFR [0.00004]<br>Other [0.0001] |
| 3-100087892-C-T | rs957967770 | p.Val514Ile | **0.562** | DCSS_PPI<br>Mutpred2_0.60 | Latino/AA [0.0001]<br>European (non-Finnish) [0.00003] |
| 3-100091455-C-T | rs770985289 | p.Ala483Thr | **0.499** | DCSS_PPI<br>Mutpred2_0.90 | European (non-Finnish) [0.00002]<br>Latino/AA [0.00002] |
| 3-100092482-A-G | rs1266227924 | p.Ile412Thr | -1.054 | DC_PPI | African/AA [0.00008] |
| 3-100091472-T-C | rs750026792 | p.Glu477Gly | -1.014 | DC_PPI | South Asian [0.000032] |
| 3-100105815-A-G | rs765666545 | p.Leu111Pro | -1.056 | near_ligsite | European (non-Finnish) [0.000008] |
| 3-100084470-C-T | rs753612125 | p.Asp589Asn | -0.534 | DCSS_PPI | African/AA[0.0000615];<br>South Asian [0.00003] |
| 3-100086973-C-T | rs775872768 | p.Gly530Ser | -0.545 | - | South Asian [0.00003] |
| 3-100093900-C-T | rs762230068 | p.Asp397Asn | -0.536 | - | African/AA [0.0001]<br>European (non-Finnish) [0.000008] |

The variant V556L has the highest predicted change in the binding affinity (ΔΔG^Affinity^ score of 0.905 Kcal/mol). V556 is a directly contacting (DC_PPI) residue at the ORF9b:TOMM70 interface. The wild type V556 in TOMM70 forms hydrophobic bonds with A68 and F69 in ORF9b, and a polar interaction with T72. The mutant V556L gains additional hydrophobic bonds with A68 and F69, resulting in increased predicted affinity for SARS-CoV-2: ORF9b.

The DCSS-variant at residue position 483 lies close to (i.e., within 5Å) the phosphorylation site in SARS-CoV-2: ORF9b i.e., S53, which is important for binding of ORF9b to TOMM70. The formation of TOMM70:ORF9b complex is regulated via phosphorylation at S53 [99]. A483 also interacts with other DC residues in TOMM70. The A483T mutation strengthens this interaction, and this mutation is predicted to be strongly pathogenic by mutpred2 (score =0.90). Figure 2 summarizes impact of all affinity-enhancing variants and their structural impact on atomic interactions at the interface.

Most affinity-increasing variants in TOMM70 are observed in the African American and American population, but with rare allele frequency i.e., less than 1% (Table 3). Most of the affinity-enhancing variants have an impact on atomic interactions of residues in the interface and are also predicted to have significant impact by various programs including CADD (score > 20), SIFT (predicted as deleterious) and predicted to be pathogenic by mutpred2 (score >0.5), as shown in Figure 2(B).

#### 2. Impact of human IFIH1 variants on SARS-CoV-2: PLPro binding

PLpro (papain-like cysteine protease) in SARS-CoV-2 is involved in a wide range of important functions, such as viral polyprotein chain processing, dysregulation of host inflammatory responses, and impairing the type I interferon (IFN-1) antiviral immune responses [100] . SARS-CoV-2: PLpro plays a key role in innate immune suppression in humans by interacting with various host substrates such as ISG15, IFIH1 and IFIT2, and others [101, 102]. Therefore, the SARS-CoV-2:PLpro is a hot spot for designing protein-protein interactor inhibitors [102].

IFIH1 (also known as MDA5) is a cytoplasmic innate immune receptor. IFIH1 is a pattern-recognition receptor which binds to viral RNAs and suppresses translation initiation [103]. IFIH1-binding to viral RNA is known to induce type I interferon response by triggering activation of antiviral immunological genes including IFN-alpha, IFN-beta and pro-inflammatory cytokines [103–105]. SARS-COV-2 employs PLpro to bind and block the activation of an IFIH1-dependent cascade of antiviral responses [104]. PLpro is suggested to bind CARD domains of IFIH1 [101, 104] and hence we modelled a complex of PLpro and IFIH1 using AlphaFold2-multimer (Figure 3).

**Figure 3:**
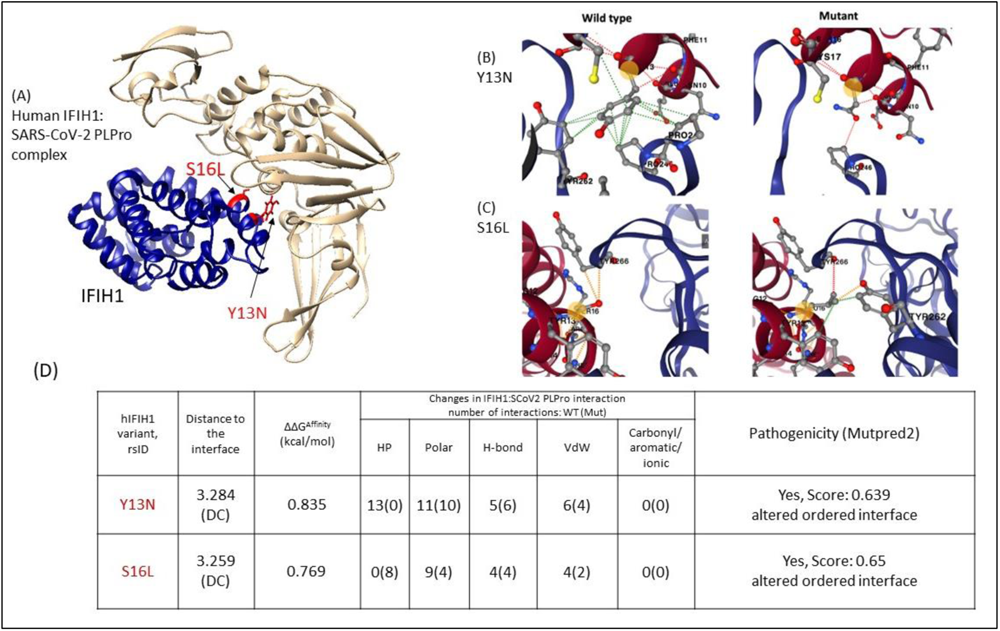
IFIH1-PLpro complex built using AlphaFold2-multimer. (A) Human IFIH1 is shown in tan and SARS-CoV-2:PLpro is shown in blue. The structural locations of affinity-enhancing variants (ΔΔG^Affinity^ ≥ 0.5 Kcal/mol) are shown in red. These include three DC-variants (S16L, Y13N) (B, C) The structural impact of DC-variants on atomic interactions at the interface, are shown. (C) Effects of affinity-enhancing variants in TOMM70 are shown in detail (figures in 2B and 2C from mCSM-PPI2). HP: Hydrophobic (green), H-bond: hydrogen bond (red), VDW: Van-der-Waals (blue), Polar (orange).

### Affinity-enhancing variants: structural impact and population distribution

Two affinity-enhancing variants in IFIH1 are predicted - both Y13N and S16L, are DC residues (Figure 3). According to gnomAD, Y13N is a common variant [allele frequency (AF) > 1%] in East Asians. This is in accordance with GenomeAsia100K, which indicates Y13N is a common variant in Northeast Asian i.e., in Japanese (AF= 0.01) and Korean populations (AF = 0.003). Upon mutation Y13N, hydrophobic interactions between aromatic rings of Y13 in wild-type IFIH1 and of P245, P246 and Y262 in viral PLpro are replaced by a stronger hydrogen bond between the N13 of the mutant IFIH1 and the P246 of PLpro (Figure 3).

The other variant significantly impacting SARS-CoV-2:PLpro binding affinity is S16L, observed to occur at a rare frequency in Europeans (non-Finnish). The side chain oxygen atom on residue S16 of IFIH1 forms two weak polar interactions with Y266 of PLpro whereas the leucine side chain in the S16L IFIH1 mutant interacts more strongly with Y266 (Figure 3b). Leucine side chain forms a hydrogen bond with viral Y266 and hydrophobic and polar interaction with Y262 (Figure 3).

#### 3. Impact of human ISG15 variants on SARS-CoV-2: PLPro binding

ISG15 (Interferon stimulated gene 15) plays a key role in the innate immune response to viral infection via a process known as ISGylation upon activation by type I interferons or by viral/bacterial infections. ISGylation (ISG15 modification) is a process whereby ISG15 protein covalently binds to other protein substrates [100, 106]. The ISGylation process acts as an antiviral defence mechanism against SARS-CoV-2 and several other RNA viruses . SARS-CoV-2 PLpro binds to ISG15 and block the ISGylation [104, 106, 107]. ISG15 binds to PLpro via LRGG motif (154-157 residues).

#### Affinity-enhancing variants: structural impact and population distribution

Two ISG15 coding variants at DC positions (S21N, L121N) are predicted to enhance the binding affinity of the ISG15-PLpro complex (Table 4, Figure 4).

**Figure 4:**
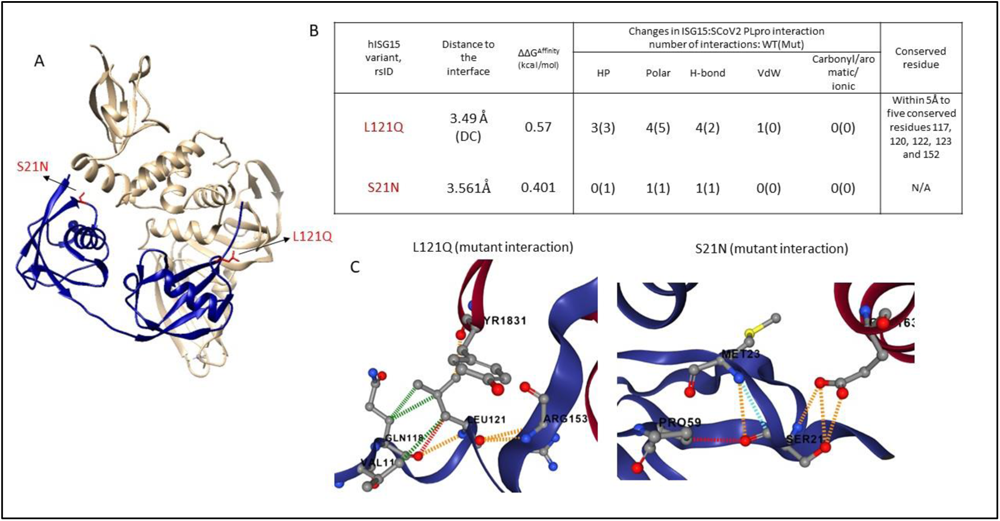
SARS-CoV-2: PLpro-human ISG15 complex (7RBS). (A) Human ISG15 (blue) in complex with SARS-CoV-2: PLpro (tan). The affinity-enhancing variants L121N and S21N are indicated in red. (B) The table summarizes the impact of affinity-enhancing mutations (wild type vs mutant) on atomic interactions at the interface. Atomic interactions associated with L121Q and S21N are shown in (C). HP: Hydrophobic (green), H-bond: hydrogen bond (red), VDW: Van-der-Waals (blue), Polar (orange)

**Table 4:** Details of variants in human ISG15 that impact binding to SARS-CoV-2:PLpro. For each variant, SNP ID, amino acid mutation, mCSM-PPI2 prediction, functional site annotations and enrichment in specific population in gnomAD are indicated. Functional sites are abbreviated as follows-DC_PPI: directly contacting interface site, DCSS_PPI, secondary shell residue, ligsite: known ligand binding site, near_ Scorecon90: proximity to conserved sites predicted using Scorecons (score ≥ *90). Variants with Mutpred2 pathogenicity score >0.50 are listed*.

| rsIDs | Mutation | McsM-PPI2 (kcal/mol) | Functional site annotations | Populations in gnomAD and [Allele frequency] |
| --- | --- | --- | --- | --- |
| rs748715915 | p.Leu121Gln | **0.57** | DC-PPI, Near_SC90 Near_LRGG motif | NFE-SWEDISH [0.00003834] |
| rs143888043 | p.Ser21Asn | **0.401** | DC-PPI | AFR [0.01648]<br>*also supported by GenomeAsia1000K<br>AMR[0.0008475],<br>NFE [0.00007794],<br>SAS[0.00009810],<br>other[0.0008333] |
| rs1477663018 | Leu85Pro | -1.002 | DCSS-PPI, Scorecons90 | NFE-North-Western [0.00002404] |
| rs1160940574 | p.Met23Thr | -1.017 | DC-PPI | NFE-North-Western [0.00002387] |
| rs761507082 | p.Arg155Trp | -1.166 | DC-PPI LRGG motif site | AMR[0.0004058]<br>ASJ[0.0001011],<br>SAS[0.00003275], |
| rs774241776 | p.Asn151Lys | -1.286 | DC-PPI | Other [0.0001647] |
|  |  |  | Near_Scorecon90 |  |
| rs750338976 | p.Arg155Gln | -2.061 | DC-PPI<br>LRGG motif site | NFE[0.00001808] |

L121Q, one of the direct contact residues in the interface, is predicted to enhance the ISG15: SCoV2 PLpro binding affinity (mCSM-PPI2 ΔΔG^Affinity^ = 0.57 kcal/mol). This variant occurs within 5Å from the key LRGG motif (the PLpro recognition site) and forms direct interaction with R153, which is adjacent to this motif, which is therefore likely to have impact on PLpro-binding. Analyses using conserved sites using Scorecons indicates that the variant lies in the structural neighbourhood (5Å) from five conserved residues (at positions 117, 120, 122, 123 and 152; with Scorecons90) of which one (W123) is also predicted to be an allosteric site (score: 0.896, predicted using Ohm [108]). This substitution is only observed in the Swedish population at rare frequency.

The second DC-variant S21N is annotated in ClinVar (ID: 475283, benign) and is associated with Mendelian susceptibility to mycobacterial diseases (also known as Immunodeficiency 38 disease)[109]. The variant is associated with severe clinical disease upon infection with weakly virulent mycobacteria (including Mycobacterium bovis and Bacille Calmette-Guerin vaccines) [110, 111]. The S21N variant is predicted to moderately increase affinity (with the borderline mCSM-PPI2 ΔΔG^Affinity^ score of 0.401 Kcal/mol)) and is a common variant (allele frequency >1%) found in the African population, as supported by multiple population databases including gnomAD [allele frequency (AF): 0.01648], GenomeAsia100K (allele frequency 0.043269) and AllofUs (allele frequency 0.014).

In the case of affinity-reducing (protective) variants (mCSM-PPI2 ΔΔG^Affinity^ ≤ -0.05 Kcal/mol), four variants (R155Q, R155Q, N151D, L85P, M23T) were predicted with significantly reduced binding affinity (ΔΔG^Affinity^<-1.0 Kcal/mol). Most of these occur only in Non-Finnish Europeans and one predominantly in the American population (R155W). Two variants (R155Q, R155W) occur within the known PLpro-recognition motif in ISG15 (LRGG motif formed by 154-157 residues) [106]. Thus, individuals carrying these mutations may be at lower risk of compromised immunity mediated by ISG15: PLpro binding.

#### 4. Impact of human IFIT2 variants on SARS-CoV-2: PLPro binding

IFIT2 (Interferon-induced protein with tetratricopeptide repeats 2) is an RNA-binding protein, and binding of RNA is known to be important for antiviral activity of IFIT2 [127]. Biochemical studies indicate that human IFIT2 binds to SARS-CoV-2:PLpro [101], however an experimental structure of the complex is not available. The modelled complex in our study, indicates SARS-CoV-2: PLpro binds at a channel-like region at the C-terminus of IFIT2, which is a known RNA-binding region (formed by K37, R184, K255, R259, R291, and K410) [128]. Three PLpro-interacting residues in IFIT2 (i.e., R259, K410 and R291) are involved in RNA-binding (depicted in Figure 5).

**Figure 5:**
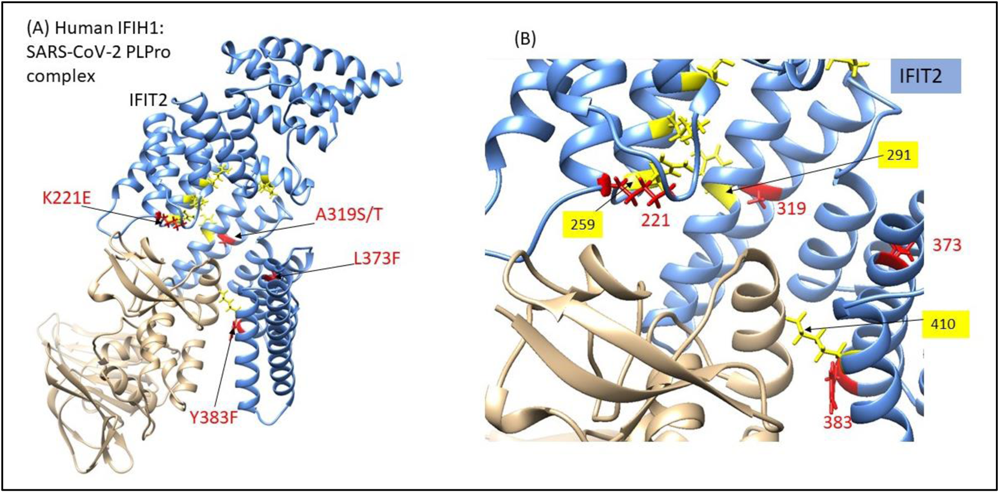
The mode of SARS-CoV-2:PLpro in complex with human:IFIT2, and mapping of affinity-enhancing variants. (A) Mapping of the affinity-enhancing variants (red) K221E, A319S/T, L373F, Y383F onto the IFIT2 (cornflower blue)-Sars-CoV-2:PLpro (tan) complex**. (B)** PLpro interacts with IFIT2 at a site which partially overlaps with a known RNA-binding region (yellow).

#### Affinity-enhancing variants: structural impact and population distribution

Four IFIT2 DCSS-variants (L373F, K221E, A319S and A319T) significantly affect the binding affinity (mCSM-PPI2 ΔΔG^Affinity^ ≥ 0.5 Kcal/mol) (see Table 5). Three of these-A319S, A319T and L373F are also predicted to be pathogenic by CADD (score ≥ 20) and SIFT scores. Two DCSS-variants in IFIT2 namely, K221E and A319(S/T) are predominant in South Asian and African/AA populations, respectively. The residues (K221E, A319S and A319T and Y383F) directly interact with residues (R259 and R291) known to be involved in the RNA-binding in IFIT2 (Figure 5B). Thus, mutations in IFIT2 protein which increase binding to SARS-CoV-2 PLpro are likely to hinder the binding of IFIT2 to RNA, and thus the normal antiviral mechanism of IFIT2.

Thus, our analyses of 3D complexes, affinity-enhancing variants, and functional sites identifies some affinity enhancing variants that could promote the binding of host immune proteins to SARS-CoV-2 proteins, thereby reducing the binding to their natural protein partners. This could mediate reduced immune responses in certain individuals.

**Table 5:**
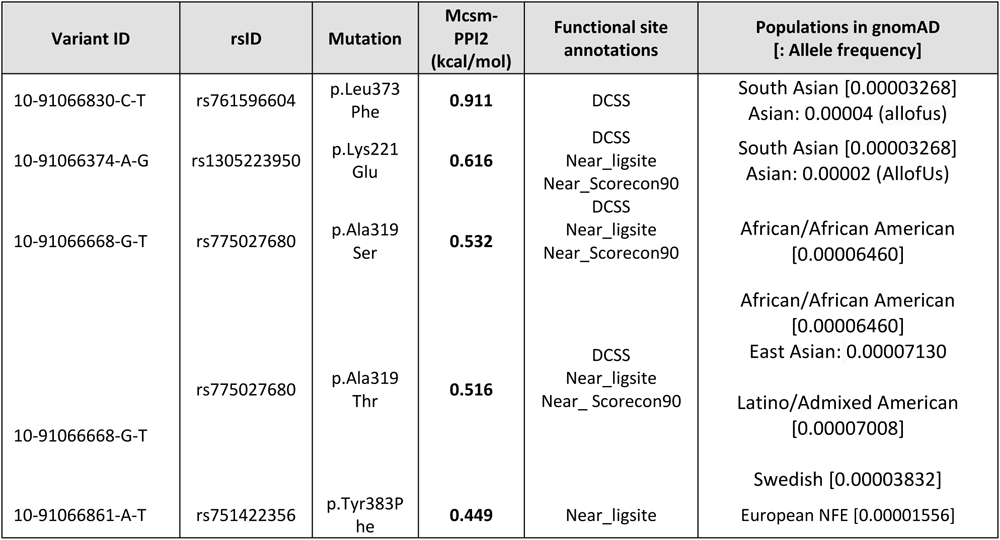
Details of affinity-enhancing variants in variants in IFIT2 (PLpro interacting protein). For each variant, SNP ID, amino acid mutation, mCSM-PPI2 prediction, functional site annotations and enrichment in specific population in gnomAD are indicated. Functional sites are abbreviated as follows-DC_PPI: means directly contacting interface site, DCSS_PPI, secondary shell residue, ligsite: indicates known ligand binding site (i.r. TNA binding site in IFIT2) and near_ligsite indicates sites that are proximal to a known ligand binding site. near_ Scorecon90: proximity to conserved sites predicted using Scorecons (score > 90).

#### Impact of variants in SARS-CoV-2 proteins

In addition to human protein variants, we also analysed the impact of viral protein variants, in Spike-RBD, Spike-NTD, PLPro, ORF9b, ORF7b, ORF6, ORF3a, NSP7, NSP15, NSP14, NSP1, N and E (using experimental/predicted complexes listed in Table 1). However, a few variants in SARS-CoV-2 proteins such as NSP14 (L6074F and N6054I), PLpro (L1774F) and Spike (S477N) are predicted to be affinity-enhancing (mCSM-PPI2 ΔΔG^Affinity^ ≥ 0.5 Kcal/mol). The impact of the variant spike-RBD S477N on enhanced ACE2-binding affinity is recently confirmed by experimental assays [37].

## Pathway enrichment analyses

We examined the biological processes and signalling pathways associated with the 14 human proteins containing affinity-enhancing variants (Table 2) using three well-established pathway enrichment databases: Gene Ontology (GO), KEGG and Reactome (Table 6). The most significantly enriched terms were related to immune functions, including viral life cycle (GO biological process) and Influenza A (KEGG). The top enriched Reactome pathway was SARS-CoV-2 Infection, confirming the association with this panel of proteins identified in earlier analyses.

**Table 6.** Top significant GO, KEGG and Reactome pathways associated with 14 SARS-CoV-2-interacting human proteins (containing affinity-enhancing variants).

| Term | Pathway | Adj. p-value |
| --- | --- | --- |
| REAC:R-HSA-9694516 | SARS-CoV-2 Infection | $2.59 \times 10^{-8}$ |
| GO:1903900 | regulation of viral life cycle | $3.19 \times 10^{-5}$ |
| KEGG:05164 | Influenza A | 0.0019 |

Since many biological processes are governed by functional modules which are highly interconnected sub-networks of protein-protein interactions, identifying functional modules containing the 14 human proteins provides a method for gaining further insights into their biological functions. This approach can also offer insights into specific mechanisms contributing to SARS-CoV-2 pathology.

To this end, we used two state-of-the-art modularity detection algorithms and applied them to two well regarded network datasets: STRING, and ConsensusPathDB. We then identified which modules contain any of the 14 genes and performed pathway enrichment analysis on each of the modules using GO (biological processes), KEGG and Reactome. The enrichment analysis was used to assign to each of the 14 genes the appropriate biological processes/pathways. For a full list of genes in each module, see supplementary file 5 and 6.

Unsurprisingly, many of the 14 genes were associated with immune response to viral infection, including interferon-inducible genes IFIT2 and IFIH1, and ISG15 involved in modulating viral replication. Induction by type I interferons (α/β), which activate other immune cells, was also associated with the transmembrane protein TRIM25.

Several symptoms, particularly in severe or long COVID-19 cases, are associated with mitochondrial dysfunction, including cytokine storms [112], which is a key pathway associated with TOMM70 associated with mitochondrion organisation. It has also been found to be involved in interferon regulation [113]. SARS-CoV-2 viral proteins replicate in the cytoplasm via translation on ribosomes, hence supporting our identification of several pathways involved in cytoplasmic or ribosomal transport, specifically NUP98, RAE1, ribosomal protein RPS3, TOMM70 and TRIM25. Other genes are involved in cell processes that are hijacked by the virus during infection. For example, ARF6 was identified as involved in exocytosis, but is used by the virus to infiltrate and infect the cell [114]. Overall, the findings of our functional analysis highlight some key mechanisms involved in the viral response to COVID-19.

### Functional families associated with small molecule inhibitors

For the 14 human proteins containing predicted affinity-enhancing variants, we analysed their associated functional families in CATH to inspect whether any of the homologous proteins in the FunFam were linked to known small molecule inhibitors. These small molecules, which may include drug-like molecules or approved drugs, were identified using the ChEMBL database (see Table 7).

**Table 7:** Human proteins with CATH-FunFams containing small molecule inhibitors. Three human proteins (IFIH1, AXL and ARF6) are observed to be linked with small molecule inhibitors from ChEMBL. The associated entries in ChEMBL are indicated. The representative 3D structure for each functional families in CATH is indicated. If the representative is not available in CATH, the representative from PDB is shown.

| Protein | CATH superfamily and functional family (CATH-FunFam) ID | Associated entry in ChEMBL | Representative 3D-structure |
| --- | --- | --- | --- |
| IFIH1 | 1.10.533.10<br>CATH-FunFam ID 77 | CHEMBL4739862<br>[CHEMBL4594258, phase 2] | 7DNI [PDB] |
| AXL | 2.60.40.10<br>CATH-FunFam ID 810 | CHEMBL4895<br>[CHEMBL3301622, approved]<br>CHEMBL4879451 | 5vxzD00 [CATH] |
| ARF6 | 3.40.50.300<br>CATH-FunFam ID 286 | CHEMBL5987<br>CHEMBL1075274, | 3lvrE04 [CATH] |

Next, we used CavityPlus to detect druggability of the protein-protein interface formed by these three proteins. CavityPlus provided strong confidence for prediction for druggability of IFIH1: PLpro interface and ARF6:NSP15-interface, and medium confidence for AXL:NTD interface (see supplementary file 7). These results are indicative of potential applications of their associated inhibitors for designing drugs targeting these protein-protein interactions.

For example, IFIH1 is associated with Selgantolimod in ChEMBL (ID: CHEMBL4594258), which is in phase 2 clinical trial. Selgantolimod is known to be a Toll Like Receptor 8 Agonist, which increases immune responses in chronic Hepatitis B patients (Reyes et al., 2019). Analyses using CavityPlus provided a strong prediction score for the presence of two cavities that occur at IFIH1-PLpro interface (the topmost cavity with score of 4177.0, Supplementary File 7). These cavities do not interfere with CARD oligomerization or its protein partner MAVS, which is required for IFIH1-induced interferon response, thus substantiating the potential use of this molecule in designing therapeutics against SARS-CoV-2 infection. Docking of this ligand with IFIH1-PLpro, provides support for binding at this cavity ([115]; see Supplementary File 8).

Likewise, cavities the interface of NTD-AXL and ARF6-NSP15 could likely be interesting targets for drug design and further experimental assays are required to substantiate their application for drug repurposing or as starting points for structure-based design of novel compounds.

## Discussion

We analysed ∼20% of SARS-CoV-2 immunity associated interactions using structural data of protein-protein complexes. Structural analyses helped in predicting affinity-enhancing variants in immunity-associated proteins and their impact on SARS-CoV-2-human protein complexes and COVID-19 susceptibility. Detection of putative affinity-enhancing variants in human proteins and information on their proximity to functional sites in the 3D structure can aid in explaining potential mechanisms associated with the suppression of normal functioning of immune proteins, thereby affecting COVID-19 susceptibility.

We applied a structural bioinformatics approach to analyse the impact of missense variants from human and viral proteins, using 19 SARS-CoV-2: human protein complexes (obtained from PDB or built using AlphaFold2-multimer/ptm). We analysed 468 coding variants in human proteins occurring at protein-protein interfaces. A total of 26 affinity-enhancing variants from 14 human proteins were predicted to significantly enhance SARS-CoV-2 binding.

A majority of these human proteins were involved in key immune pathways and associated with antiviral activity against SARS-CoV-2, including Interferon stimulating genes (ISG15, IFIT2); important receptors (such as IFIH1, TOMM70); proteins involved in nucleocytoplasmic shuttling of viral mRNA (NUP98 and RAE1), proteins involved in cellular translation machinery (RPS2 and RPS3) and cell entry receptors via spike-binding (ACE2, KREMEN1 and AXL). Among these 14 proteins, experimental assays have been performed on spike-ACE2, substantiating the role of the predicted spike affinity-enhancing variants in COVID-19 susceptibility and transmission [37, 38], while variants in the remaining proteins are reported for the first time in this study.

The modelling of complexes using AlphaFold2-multimer/ptm helps to provide structural insights into the mechanisms of SARS-CoV-2 binding and increased the structural coverage of the complexes. We propose that affinity-enhancing variants in key-immunity associated human proteins could promote their binding to SARS-CoV-2 proteins, competing with protein partners or substrates in immune pathways, and this in turn, may have an impact on COVID-19 susceptibility. This finding is in line with previous experimental studies on SARS-CoV-2: human interactions that affect natural immune pathways [116, 117]. For example. Li et al. [117] suggest that binding of SARS-CoV-2: ORF6 to human: NUP98-RAE1 complex competitively inhibits mRNA binding (to NUP98-RAE1), which is essential for its immune function. Likewise, overexpression of SARS-CoV-2: Nucleoprotein protein is observed to be associated with the attenuation of RIG-I-mediated interferon production via binding to TRIM25 and thus interrupting the interaction between TRIM25 and RIG-I (which is its natural protein partner) [116].

The SARS-CoV-2: human protein interactions associated with these 14 human proteins could be attractive targets for drug design that target the protein-protein interfaces. In particular, SARS-CoV-2: PLpro is known to be a promising target for designing protein-protein interaction inhibitors [118]. In our analyses, we modelled PLpro interactions with ISG15, IFIH1 and IFIT2. Interestingly, PLpro interactor protein IFIH1 is associated with drug-associated functional family in CATH. Likewise, we observed druggable CATH-FunFam for AXL and ARF6 proteins (interacting with Spike-NTD and NSP15 in SARS-CoV-2, respectively), where drugs have previously been identified that bind to structurally similar paralogs. The domain relatives within CATH-FunFams exhibit highly conserved drug binding sites and have the potential to be the druggable entities within drug targets, as shown in [60]. Thus, further studies targeting such immune proteins and PLpro-mediated interactions would be helpful.

We suggest monitoring both common and rare variants in human proteins, that are predicted to cause significant impact and thus likely to be associated with disease pathogenicity or susceptibility. Two affinity-enhancing variants – one from IFIH1 (Y13N) and one from ISG15 (S21N), are observed to occur at > 1% allele frequency in East Asians and African population, respectively. Most of the variants identified in our study have allele frequencies (<1%). A growing number of studies support the key role of rare variants in causing susceptibility/severity to COVID-19 [12, 26–32].

We also provide a catalogue of protective variants from 14 proteins, particularly in ISG15 and TOMM70. These variants are observed to significantly reduce binding affinities of the human proteins to their SARS-CoV-2 partners. Earlier data showed a good correlation between predicted affinity-reducing variants and experimental observations [38]. Thus, affinity-reducing variants reported in this study could provide an explanation for why some individuals in specific populations are less likely to experience SARS-CoV-2 associated immune evasion.

Whilst we observed only two common variants in certain specific ethnic groups, occurrence of certain rare affinity-enhancing variants could also lead to increased susceptibility in individuals carrying them. Though the data suggest some role of genetic variation in COVID-19 susceptibility, the role of social and environmental factors should also be studied. In addition, our study is based on on computational prediction of changes in the binding affinity. Experimental data is available for variants in only a limited number of complexes such as spike-ACE2 and ORF9b-TOMM70. Future experimental studies would be helpful for proposed affinity-enhancing mutations in other immune proteins.

## Conclusions

We used structural bioinformatics approaches to predict human and viral protein variants affecting COVID-19 susceptibility and to suggest repurposing of therapeutics based on CATH functional family data associated with small molecules/drugs. A total of 26 affinity-enhancing variants are reported in our study and we discuss their structural impact in the context of functional sites using 3D structures of SARS-CoV-2: human complexes. The protocol designed in this study could be extended to analyse other protein interactions as more structures are experimentally determined and more powerful tools for protein structure prediction emerge. Our approach could be helpful in future studies of not only for COVID-19 but also other emerging infectious diseases.

## Data availability

The data supporting the conclusions of the study is made available in the Supplementary files 1 to 8. The dataset of SARS-CoV-2:Human 3D-complexes used in this study is provided in Supplementary file 9.

### Author Contributions

CO: the conception of the work; VW,PA,NS,NB: data acquisition, VW,SD,NS, LW,YG,JW: analysis, VW,CO,PA,YG,LW, SD: interpretation of data; PA: designed FunVar pipeline used in the work. VW,CO,PA,YG,LW,VW,SD,IS: drafted the work. All authors have approved the submitted version. All authors have agreed to be accountable for their contributions in this work.

## Conflicts of interest

Authors declare that there are no Conflicts of interest.

## Funding

VPW acknowledges funding from UKRI BBSRC grant BB/W003368/1 ; PA: Wellcome Trust grant 221327/Z/20/Z. NB: Wellcome Trust grant 221327/Z/20/Z; NS - BB/S020144/1; LW - BB/M009513/1; S.D.L. is funded by a Fundamental Research Grant Scheme from the Ministry of Higher Education Malaysia [FRGS/1/2020/STG01/UKM/02/3]; Ian Sillitoe is funded by BBSRC [BB/R014892/1]; YG acknowledges funding from BBSRC [60175814].

## Acknowledgements

David Gregory for setting up Colabfold at UCL cluster. Dr. Stuart MacGowan for his comments on the analyses. Molecular graphics and analyses performed with UCSF Chimera, developed by the Resource for Biocomputing, Visualization, and Informatics at the University of California, San Francisco, with support from NIH P41-GM103311.

## Notes

### Competing Interest Statement

The authors have declared no competing interest.

