## Supplementary material for "Predicting human and viral protein variants affecting COVID-19 susceptibility and repurposing therapeutics": Supplementary file 4.pdf

**Supplementary file 4 : Analysis of affinity-enhancing variants in human proteins**

#### I. Table 1: Human proteins containing affinity-enhancing variants and their analyses in the context of functional sites

| Human protein | The position of affinity-enhancing mutation to the functional site (interface, predicted conserved site (Scorecons90) or known ligand/substrate binding site |
|---|---|
| **ACE2** | - G326E is a secondary shell (DCSS) residue to the interface.<br>- V447F is a conserved residue identified using CATH-FunFams (Scorecons90). |
| **KREMEN1** | - V191I is a direct contact (DC) residue and within 5Å of three conserved residues.<br>- Y68H is a secondary shell and conserved residue, and in a cluster of other conserved residues. |
| **AXL** | - V38M is a direct contact residue. |
| **TOM70** | - V556 is a direct contact residue.<br>- L576 is a secondary shell residue to the interface.<br>- A591 is a direct contact residue. |
| **RPS3:NSP1** | - V91 is within 5Å of one conserved and probable allosteric site. |
| **PALS1:E protein** | - L484 is a conserved residue, and in a cluster of other conserved residues.<br>- L321 is a secondary shell residue to the interface. L321F is predicted to pathogenetic impact on the protein function. |
| **TRIM25:N protein** | - A466 is a secondary shell residue to the interface and conserved residue, and in a cluster of other conserved residues. A466T is predicted to pathogenetic impact on the protein function. |
| **ARF6:NSP15** | - L166 is a conserved residue, and in a cluster of other conserved and predicted allosteric residues. L166F is predicted to pathogenetic impact on the protein function |
| **TRIMM:NSP14** | - I105 is a secondary shell residue to the interface and predicted allosteric site, and in a cluster of other predicted allosteric sites. |
| **ISG15: PLpro** | - L121 is a direct contact residue and within 5Å of one conserved and probable allosteric site. |
| **IFIH1: PLpro** | - Y13 is a direct contact residue. Y13N is predicted to pathogenetic impact on the protein function.<br>- S16 is a direct contact residue. S16L is predicted to pathogenetic impact on the protein function. |
| **IFIT2:PLpro** | - L373 is a secondary shell residue to the interface and within 5Å of three conserved residues.<br>- K221 is a secondary shell residue to the interface and within 5Å of three conserved residues.<br>- A319 is and within 5Å of three conserved residues. |
| **hNUP98-hRAE1:ORF6** | - T190S is Scorecons90 site, part of Gle2-binding sequence (GLEBS) motif of Nup98 (residues 157–213) |

#### II. Structure analyses of affinity-enhancing variants: Changes in atomic interactions

**Table 2: Impact of affinity-enhancing variant in AXL**

| hAXL variant, rsID | Distance to the interface (Å) | $\Delta\Delta G^{Affinity}$ (kcal/mol) | Changes in hAXL:SCoV2 NTD interaction number of interactions: WT (Mut) | | | | | Conserved residue and/or within 5Å of conserved residues | Grantham score | Allosteric site (OHM) | Pathogenic (MutPred2) | Max population and allele frequency in gnomAD |
|---|---|---|---|---|---|---|---|---|---|---|---|---|
| | | | HP | Polar | H-bond | VdW | Carbonyl/ aromatic/ | | | | | |
| V38M rs781049505 | 4.0 (DC) | 0.409 | 4(1) | 1(3) | 0(0) | 0(0) | 0(0) | No | 21 (conservative) | No | No | Latino/Admixed Americans (0.00006324)<br>North-Western Europeans (0.00007390)<br>Jmorp - Japanese (0.00004) |

**Table 3: Impact of affinity-enhancing variants in RPS3**

| hRPS3 variant, | Distance to the | $\Delta\Delta G^{Affinity}$ (kcal/mol) | Changes in hRPS3:SCoV2 nsp1 interaction number of interactions: WT (Mut) | Conserved residue and/or | Grantham score | Allosteric site | Pathogenic (MutPred2) | Max population and allele |
|---|---|---|---|---|---|---|---|---|

| | | | HP | Polar | H-bond | VdW | Carbonyl/aromatic/ionic | | | | | frequency in gnomAD (unless stated otherwise) |
|---|---|---|---|---|---|---|---|---|---|---|---|---|
| V91I rs143925312 | 27.97 | 0.611 | 1(2) | 1(2) | 1(2) | (0)1 | 0(0) | No but within 5 Å of a conserved residue (R94, sc: 0.979) | 29 (conservative) | No but within 5 Å of an allosteric residue (R94, aci score: 0.865) | No | Koreans (0.005762)<br><br>GenomeAsia100k (0.002) |

**Table 4: Impact of affinity-enhancing variants in Kremen1**

| hKREMEN1 variant, rsID | Distance to the interface (Å) | $\Delta\Delta G^{Affinity}$ (kcal/mol) | Changes in hKREMEN1:SCoV2 RBD interaction number of interactions: WT(Mut) | | | | | Conserved residue and/or within 5Å of conserved residues | Grantham score | Allosteric site (OHM) | Pathogenic (MutPred2) | Max population and allele frequency in gnomAD (unless stated otherwise) |
|---|---|---|---|---|---|---|---|---|---|---|---|---|
| | | | HP | Polar | H-bond | VdW | Carbonyl/ aromatic/ | | | | | |

| | | | | | | | | | | | | |
|---|---|---|---|---|---|---|---|---|---|---|---|---|
| V191I rs753351748 | 3.25 (DCEX) | 0.9 | 2(6) | 3(6) | 2(0) | 1(1) | 0(0) | No but within 5Å of three conserved residues (92, 93 and 110) | 29 (conservative) | No | No | South Asians: 0.00009803<br><br>Latino/Admixed Americans: 0.00005793<br><br>Swedish: 0.00003831<br><br>Indigenome: 0.0015 |
| Y68H rs532050281 | 3.77 (DCEX) | 0.65 | 9(4) | 6(3) | 2(0) | 0(4) | ionic 0(1) | Yes (sc: 0.977) and within 5Å of three conserved residues (60, 67, 89, 90 and 98) | 83 (moderately conservative) | No | No | Southern Europeans: 0.0005259<br><br>Swedish: 0.00003830<br><br>Latino/Admixed Americans: 0.00002897 |

**Table 5: Impact of affinity-enhancing variants in PALS1**

| hPALS1 variant, rsID | Distance to the interface (Å) | $\Delta\Delta G^{Affinity}$ (kcal/mol) | Changes in hPALS1:SCoV2-E interaction number of interactions: WT (Mut) | | | | | Conserved residue and/or within 5Å of conserved | Grantham score | Allosteric site (OHM) | Pathogenic (MutPred2) | Max population and allele frequency in gnomAD (unless stated otherwise) |
|---|---|---|---|---|---|---|---|---|---|---|---|---|
| | | | HP | Polar | H-bond | VdW | Carbonyl/aromatic/ | | | | | |
| L484F rs372266455 | 24.0 | 0.738 | 4(15) | 1(1) | 0(0) | 0(0) | 0(0) | (sc: 1) and within 5 Å of conserved resid | 22 (conservative) | No | No | North-Western Europeans: 0.0001182<br><br>Southern European: |

| | | | | | | | | | | | | |
|---|---|---|---|---|---|---|---|---|---|---|---|---|
| | | | | | | | | ues: 482, 486, 496, 577, 655 | | | | 0.00008618<br><br>Latin/Admixed American: 0.00002827<br><br>Europeans: 0.00025 (allofus) |
| L321F rs749254713 | 3.65 (DCEX) | 0.635 | 5(10) | 5(5) | 0(3) | 2(0) | 0(2) | No | 22 (conservative) | No | (score = 0.625) Altered Metal binding | Ashkenazi Jewish: 0.0001256 |

**Table 6: Impact of affinity-enhancing variants in TRIM25**

| hTRIM25 variant, rsID | Distance to the interface (Å) | $\Delta\Delta G^{Affinity}$ (kcal/mol) | Changes in hTRIM25:SCoV2-N interaction number of interactions: WT (Mut) | | | | | Conserved residue and/or within 5Å of conserved residues | Grantham score | Allosteric site (OHM) | Pathogenic (MutPred2) | Max population and allele frequency in [illegible] |
|---|---|---|---|---|---|---|---|---|---|---|---|---|
| | | | HP | Polar | H-bond | VdW | Carbonyl/aromatic/ | | | | | |
| A466T rs141649169 | 7.0 (DCEX) | 0.593 | 7(10) | 5(5) | 0(5) | 0(2) | 0(0) | sc: 0.987, within 5Å of 9 conserved residues: 462, 465, 472, 488, 494, 495, 500, 501 and 502. | 58 (moderately conservativ) | No | score: 0.733, leads to altered metal binding | Africans/African Americans: 0.0002676<br><br>GenomeAsia100k (Mongolia): 0.0014 |

**Table 7: Impact of affinity-enhancing variants in TRIMM**

| hTRIMM variant | Distance to the interface (Å) | $\Delta\Delta G^{Affinity}$ (kcal/mol) | Changes in hTRIMM:SCoV2 nsp14 interaction number of interactions: WT (Mut) | | | | | Conserved residue and/or within 5Å of conserved residues | Grantham score | Allosteric site (OHM) | Pathogenic (MutPred2) | Max population and allele frequency in [illegible] |
|---|---|---|---|---|---|---|---|---|---|---|---|---|
| | | | HP | Polar | H-bond | VdW | Carbonyl/ aromatic/ | | | | | |
| I105F | 9.4 (DCEX) | 0.551 | 18 (31) | 7(10) | 2 (5) | 1(2) | aromatic 0(11) | No | 21 (conservative) | Yes (aci score: 0.93), and 5Å of 16 allosteric sites (residues 95 to 110) | No | Southern European: 0.00008731 |

**Table 8: Impact of affinity-enhancing variants in ARF6**

| hARF6 variant, rsID | Distance to the [illegible] | $\Delta\Delta G^{Affinity}$ (kcal/mol) | Changes in hARF6:SCoV2 nsp15 interaction number of interactions: WT (Mut) | | | | | Conserved residue and/or within 5Å of conserved residues | Grantham score | Allosteric site (OHM) | Pathogenic (MutPred2) | Max population and allele frequency in gnomAD |
|---|---|---|---|---|---|---|---|---|---|---|---|---|
| | | | HP | Polar | H-bond | VdW | Carbonyl/ aromatic/ | | | | | |
| L166F rs748170831 | 18 | 0.713 | 14(19) | 6(5) | 3(6) | 0(4) | aromatic 0(14) | sc: 0.969, and within 5 Å of 14 conserved residues: 33, 52, 54, 59, 61, 118, 120, 152, 161, 162, 164, 166, 168 and 169 | 22 (conservative) | aci score: 0.903, and within 5 Å of 6 predicted allosteric residues: 164 to 169 | No | East Asian: 0.00005447<br>IndiGenomes: 0.0005 |

**Table 9: Impact of affinity-enhancing variants in ACE2**

| hACE2 variant, rsID | Distance to the interface (Å) | SCoV2 VOC | ΔΔG[Affinity] (kcal/mol) by mCSM-PPI2 | Changes in hACE2:SCoV2 RBD interaction number of interactions: WT(Mut) | | | | | Conserved residue and/or within 5Å of conserved residues (Scorecons) | Grantham score | Allosteric site (OHM) | Pathogenic (MutPred2) | Max population and allele frequency in gnomAD |
|---|---|---|---|---|---|---|---|---|---|---|---|---|---|
| | | | | HP | Polar | H-bond | VdW | Carbonyl/ aromatic/ | | | | | |
| V447F rs776328956 | 40.0 | γ | 0.892 | 4(24) | 3(4) | 4(4) | 1(0) | aromatic 0(6) | Conserved (sc: 0.963), and within 5Å of 14 conserved residues: 236, 237, 240, 442 - 451, 584 and 588. | 50 (conservative) | No | No | Finnish: 0.0002712 Swedish: 0.0003146 |
| | | β | 0.794 | 9(21) | 2(5) | 2(3) | 1(3) | aromatic 0(2) | | | | | |
| | | α | 0.743 | 9(27) | 5(7) | 2(3) | 0(2) | Carbonyl 1(1) | | | | | |
| | | WT | 0.657 | 12(18) | 2(7) | 1(3) | 0(5) | Carbonyl 0(1) | | | | | |
| | | δ | 0.566 | 6(20) | 3(5) | 4(4) | 0(2) | 0(0) | | | | | |
| G326E rs759579097 | 5.0 (DCEX) | WT | 0.718 | 0(0) | 7(12) | 0(0) | 0(1) | Carbonyl 1(0) | No but within 5Å of 4 conserved residues: 327, 328, 330 and 331. | 98 (moderately conservative) | No | No | Africans/ African Americans: 0.0001056 |

II. **Functional families associated with human proteins containing affinity-enhancing variants**

| Human protein | Functional Family | DOP score |
|---|---|---|

| | | |
|---|---|---|
| **KREMEN1** | 2.40.20.10 | 79.2 |
| **AXL** | 2.60.40.10 | 80.1 |
| **TOM70** | 1.25.40.10 | 98.4 |
| **RPS3** | 3.30.1140.32 | 91.4 |
| **PALS1** | 3.40.50.300 | 81.7 |
| **TRIM25** | 3.30.40.10/45 | 91.9 |
| **ARF6** | 3.40.50.300 | 73.4 |
| **TRIMM** | 2.60.120.920/6 | 82.0 |
| **ISG15** | 3.10.20.90/240 | 81.9 |
| **IFIH1** | 1.20.1320.30 | 73.9 |
| **IFIT2** | 1.25.40.10/32 | 98.4 |
| **NUP98** | 1.10.10.2360 /1 | 96.7 |

**Table 10.** The human proteins and their corresponding CATH Functional Family and DOP score. DOPS calculates the degree of diversity in corresponding functional family's MSA based on the different conservation scores and frequencies DOP score is a value between 0 (for zero diversity) and 100 (for high diversity). Only MSAs with a DOPs score over 70 were considered for further analyses.

### Figures:

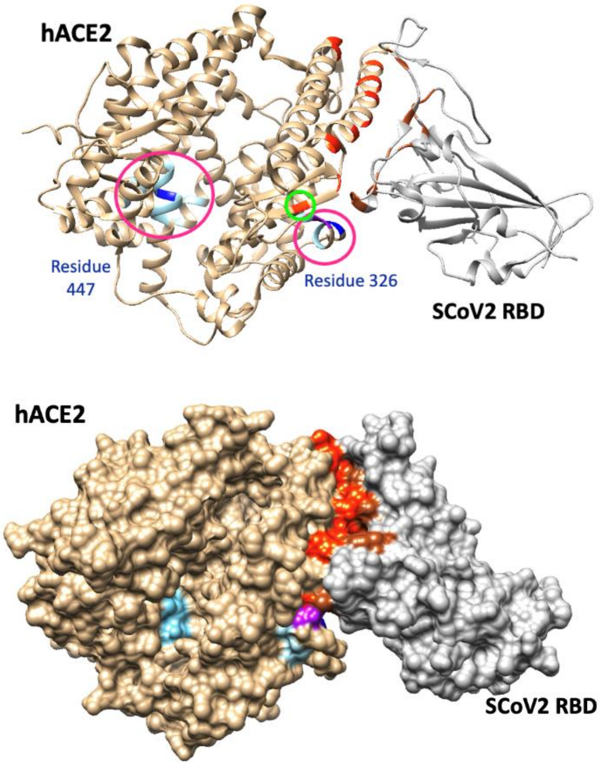


Figure S4_1. Ribbon (left) and space-filled (right) models of hACE2:SCoV2 RBD complex. The positions of two affinity-enhancing residues: 447 and 326 are shown in dark blue and conserved residues in their vicinity in light blue. The area is marked in a pink circle. Asn330 (in purple) is one of the conserved residues within 5Å of a direct contact residue: 357 (in a green circle). The red and brown residues are the directcontact residues in hACE2 (tan) and SCoV-\2 RBD (grey), respectively.

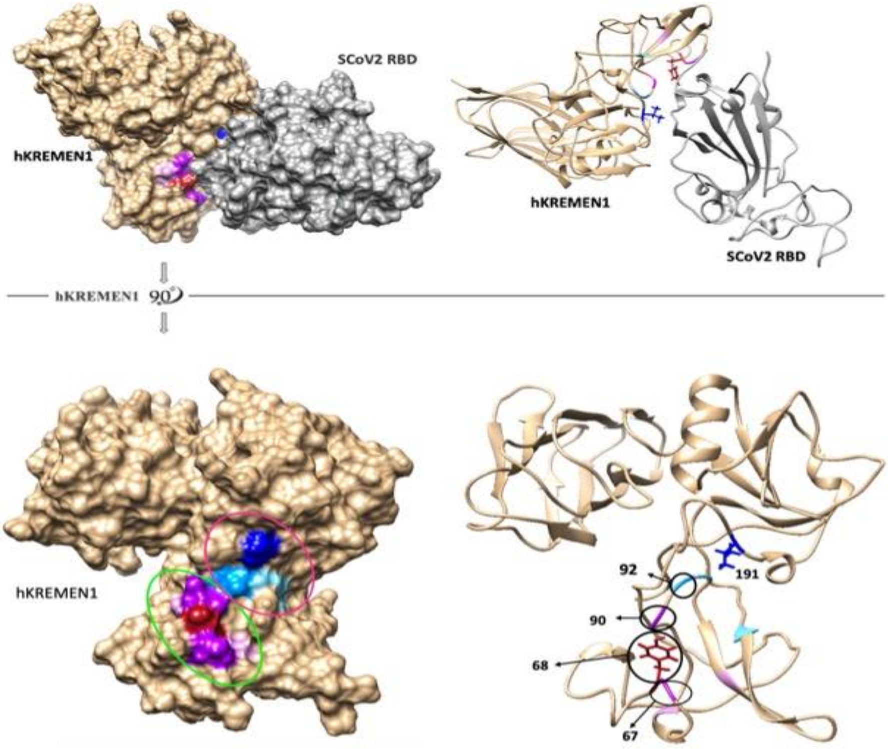


Figure S4_”. Positions of hKREMEN1 affinity-enhancing residues: 191 and 68. Top) The space-filled (left) and Ribbon (right) models of hKREMEN1:SCoV2 RBD complex. Bottom-left) A 90-degree rotation of hKREMEN1 illustrates the position of 191 (navy) and 68 (brown), their neighbouring conserved residues in red and green ovals in the space-filled model. Bottom-right) this figure presents the hKREMEN1 ribbon model with colour-coded residues for position 92 (cyan) which is 5Å from position 191 and a direct contact residue interacting with SCoV-2 RBD. Positions 67 and 90 are 5Å from position 68 and direct contact residues interacting with SCoV-2 RBD. The unlabelled blue and pink residues are the other conserved residue 5Å from positions 191 and 68, respectively. hKREMEN1 (tan) and SCoV-2 RBD (grey).

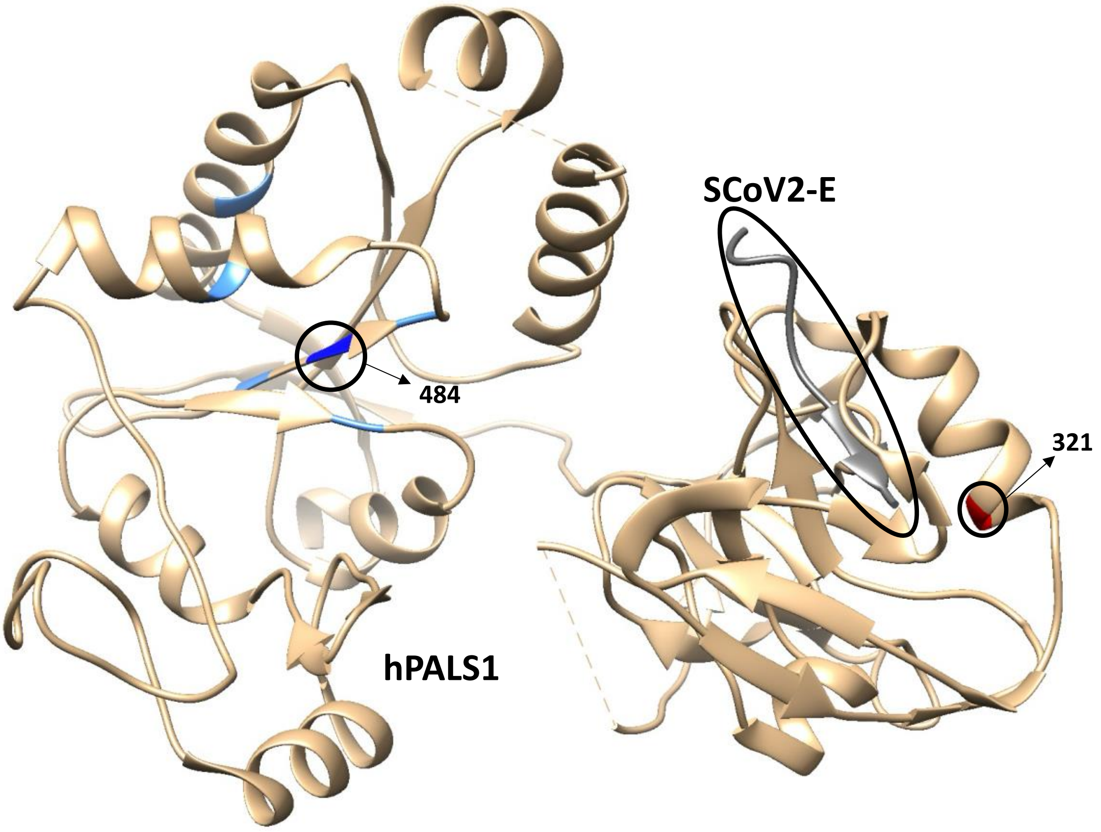


Figure S4_3. Ribbon models hPALS1: SCoV2-E complex. The position of affinity-enhancing residues: 484 is shown in dark blue and the four conserved residues within 5Å in light blue. Another affinity-enhancing residue: 321 (red) is within 5Å of direct contact residues. hPALS1 (tan) and SCoV-2-E (grey).

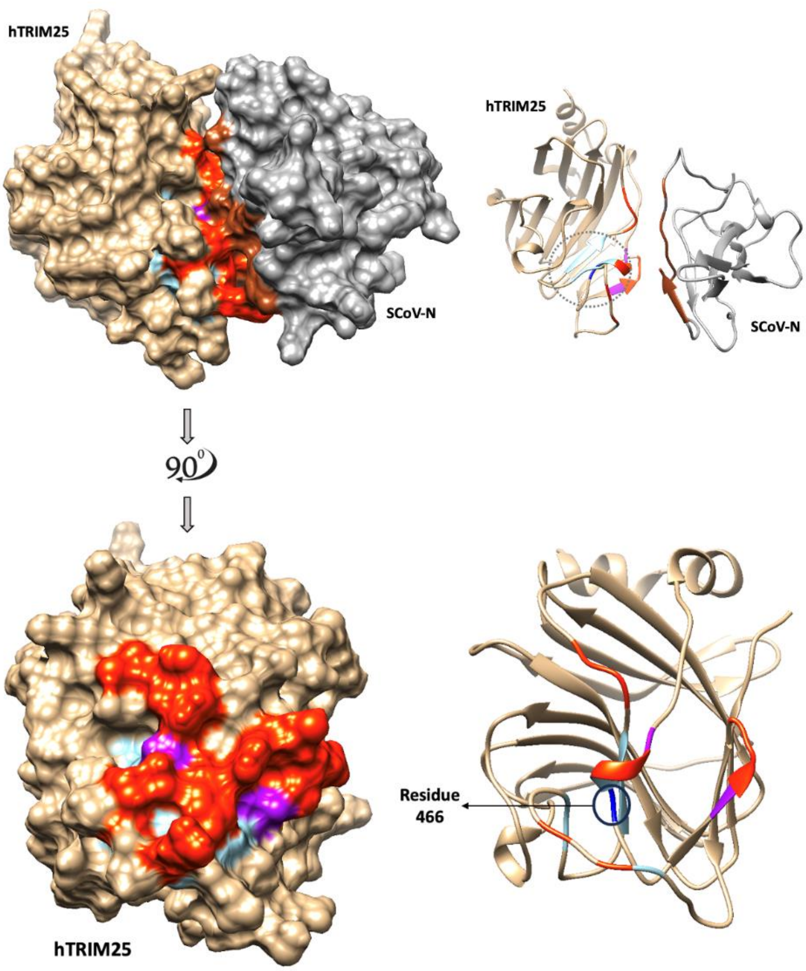


Figure S4_4: Position of hTRIM25 affinity-enhancing residues: 466. Top) The space-filled (left) and Ribbon (right) models of hTRIM25:SCoV2-N complex. Bottom) A 90-degree rotation of hTRIM25 illustrates the position of 466, its vicinity conserved residues, and the interface residues. Position 466 is shown in dark blue and its conserved residues within 5Å in light blue. The purple residues: 462 and 472 of which are in the vicinity of conserved residues are also direct contact residues. The area is marked in a dashed black circle in the top figure. The red and brown residues are the direct contact residues in hTRIM25 (tan) and SCoV-2-N (grey), respectively.

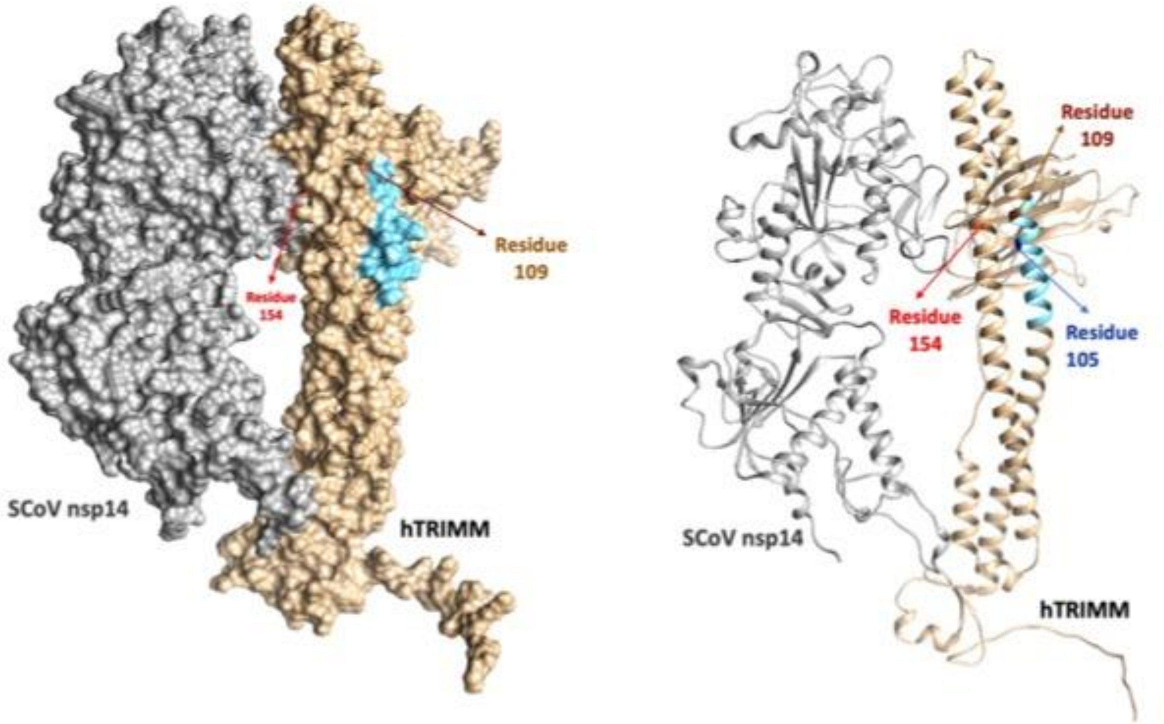


Figure S4_5. The position of residue 105 and predicted fifteen allosteric residues within 5Å of residue 105 in space-filled (left) and Ribbon (right) models in hTRIMM:SCoV2 NSP14 complex. The position 105 is in dark blue and its neighbouring allosteric positions in light blue. One of the predicted allosteric sites, position 109 in brown, is within 5Å of a direct contact residue (position 154, in red). hTRIMM (tan) and SCoV-2 nsp15 (grey).

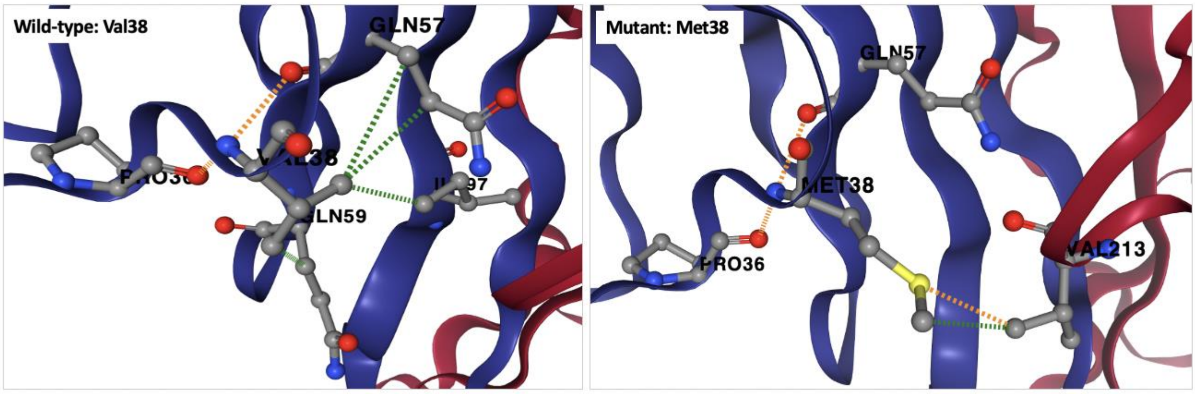


Figure S4_6. Impact of Val38Met in AXL:SCoV2 NTD affinity. The hydrophobic side chain of Met38 in AXL (blue ribbon) has hydrophobic (green dashed line) and polar (orange dashed line) interactions with Val213 of SCoV2 NTD (red ribbon) which leads to enhanced binding affinity between AXL and SCoV2 NTD.

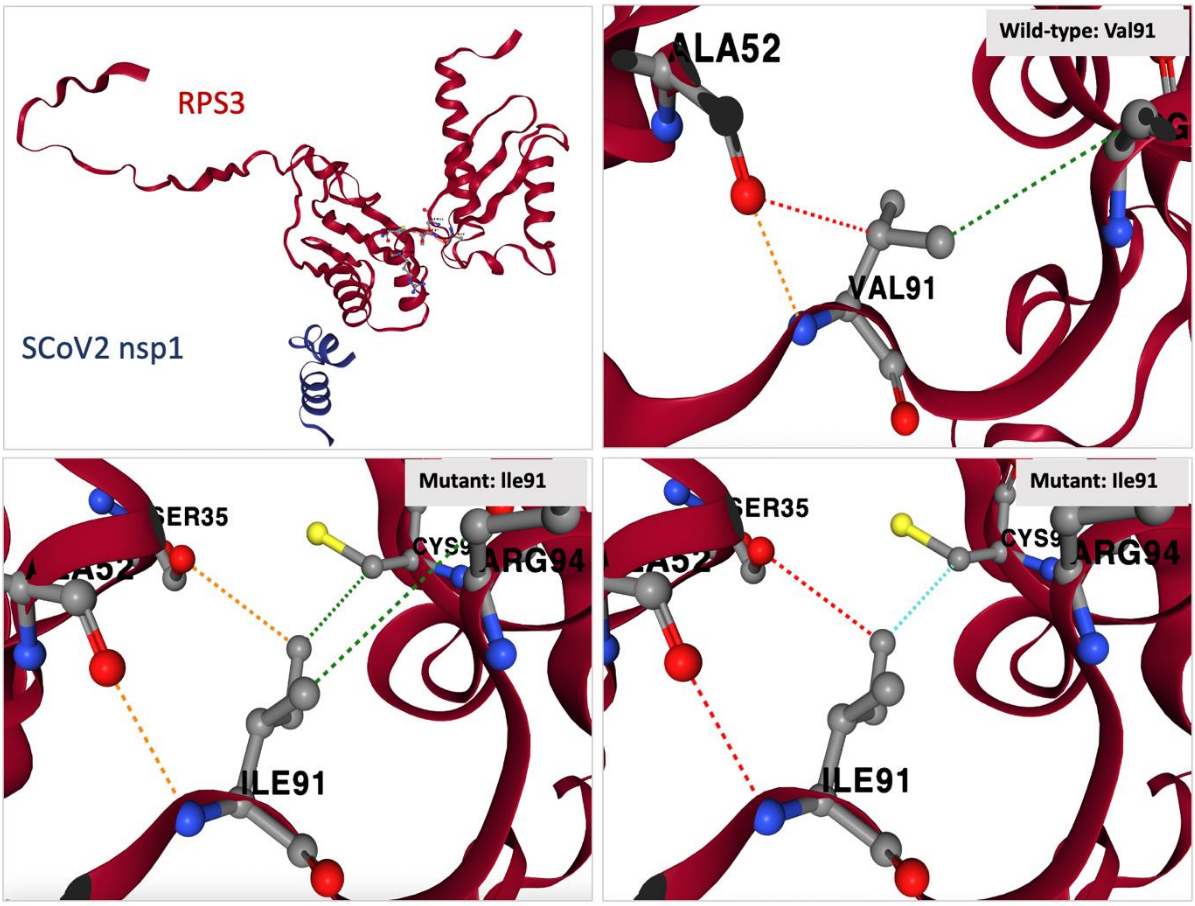


Figure S4_7. Position of RPS3 residue 91 and impact of mutation Val91Ile in RPS3:SCoV2 nsp1 complex. RPS3 and SCoV2 nsp1 are shown in red and blue, respectively. The interaction between Val91 (wild-type) and Ile91 (mutant) are shown as hydrophobic: green dash lines, Van der Waals: cyan dash lines, polar: orange dash lines and, hydrogen bond: red dash lines.

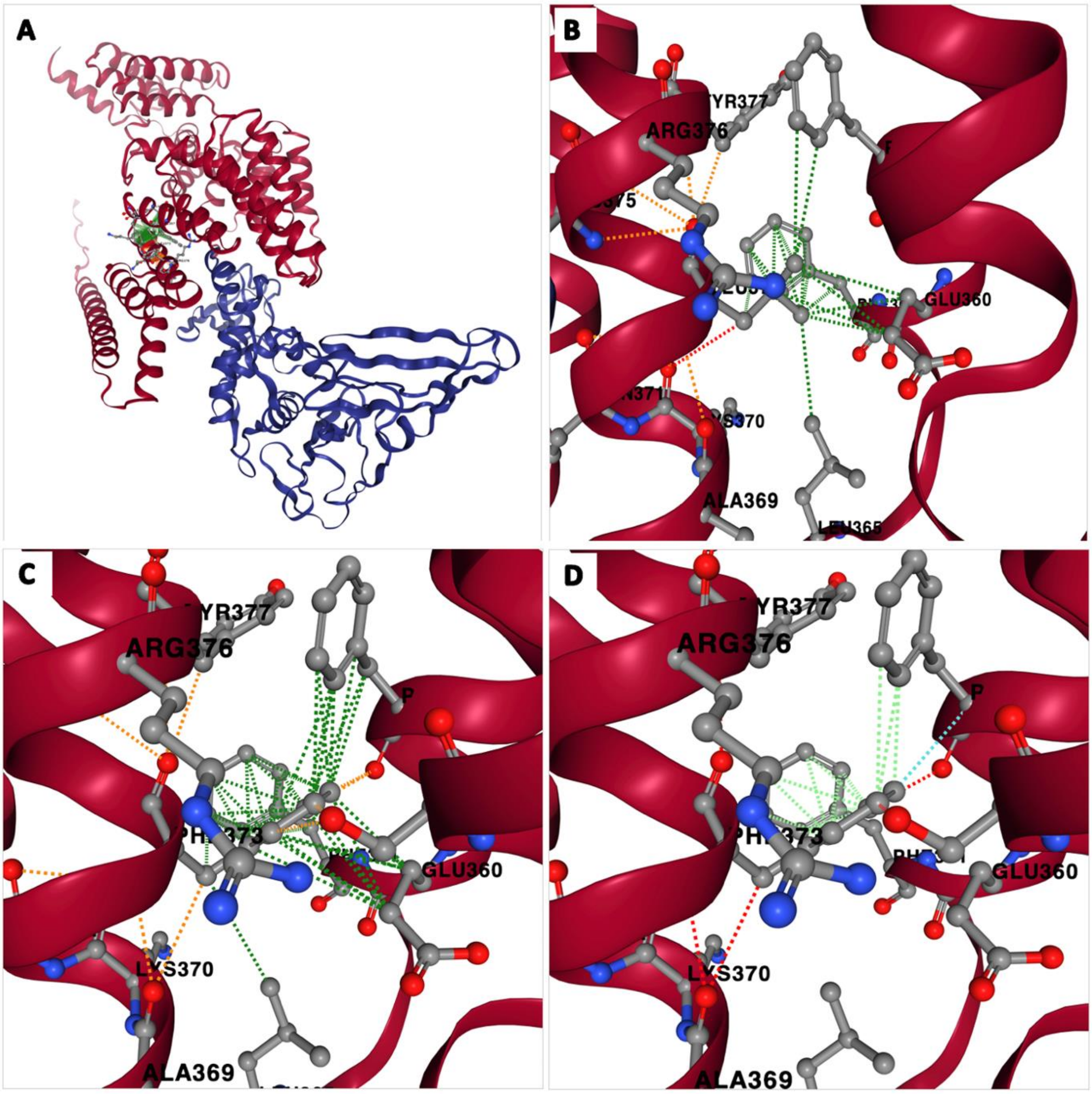


Figure S4_8. Position of residue 373 in IFIT2 and impact of mutation Leu373Phe in IFIT2:SARS-CoV-2 PLpro complex. A) Ribbon model IFIT2:SARS-CoV-2 PLpro complex and the location of residue 373. IFIT2 and SARS-CoV-2 PLpro are shown in red and blue, respectively. B) Illustration of polar and hydrophobic bonds between Leu373 and the neighbouring residues. C, D) Impact of Leu373Phe on the vicinity residues and formation of new hydrogen bonds and Van der Waals contacts. Hydrophobic (green dash lines), aromatic (light green dash), Van der Waals (cyan dash lines), polar (orange dash lines) and, hydrogen (red dash lines).

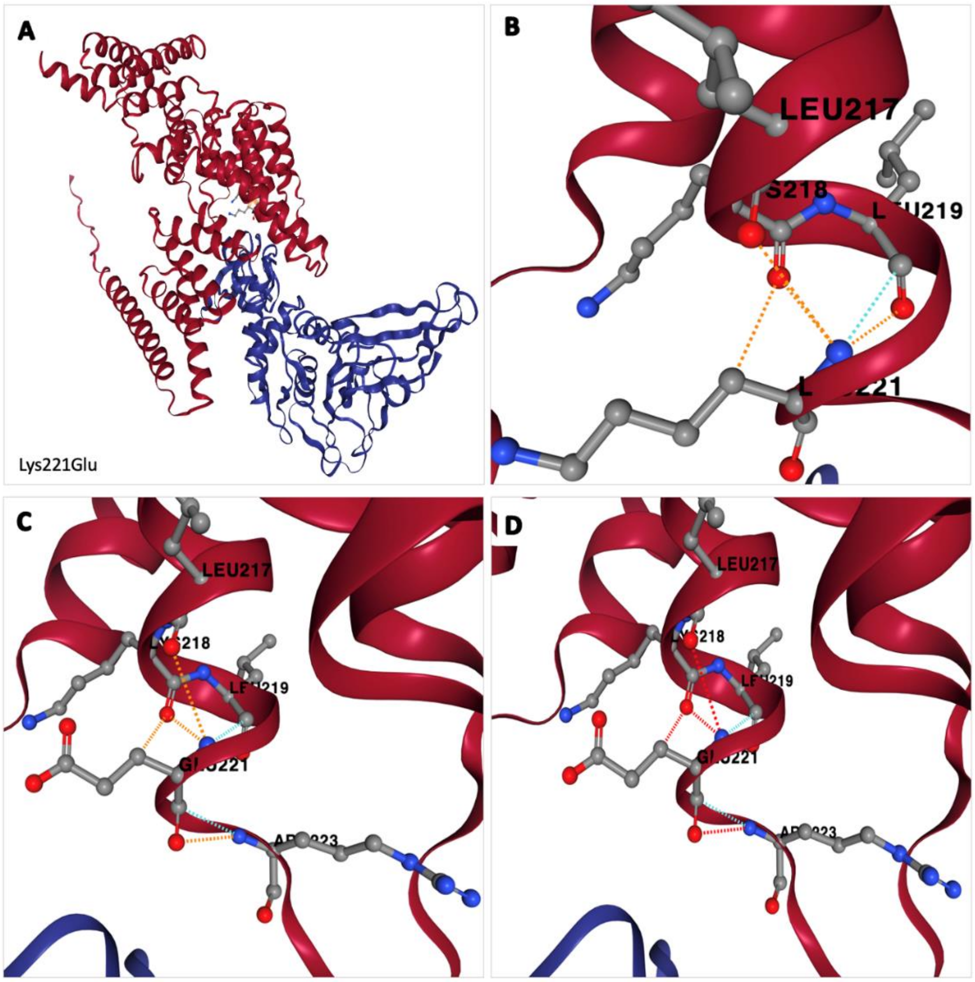


Figure S4_9. Position of residue 221 in IFIT2 and impact of mutation Lys221Glu in IFIT2:SARS-CoV-2 PLpro complex. A) Ribbon model IFIT2:SARS-CoV-2 PLpro complex and the location of residue 221. IFIT2 and SARS-CoV-2 PLpro proteins are shown in red and blue, respectively. B) Illustration of polar and hydrophobic bonds between Lys221 and the neighbouring residues. C, D) Impact of Lys221Glu on the vicinity residues and formation of new hydrogen bonds and Van der Waals contacts. Van der Waals (cyan dash lines), polar (orange dash lines) and, hydrogen (red dash lines).

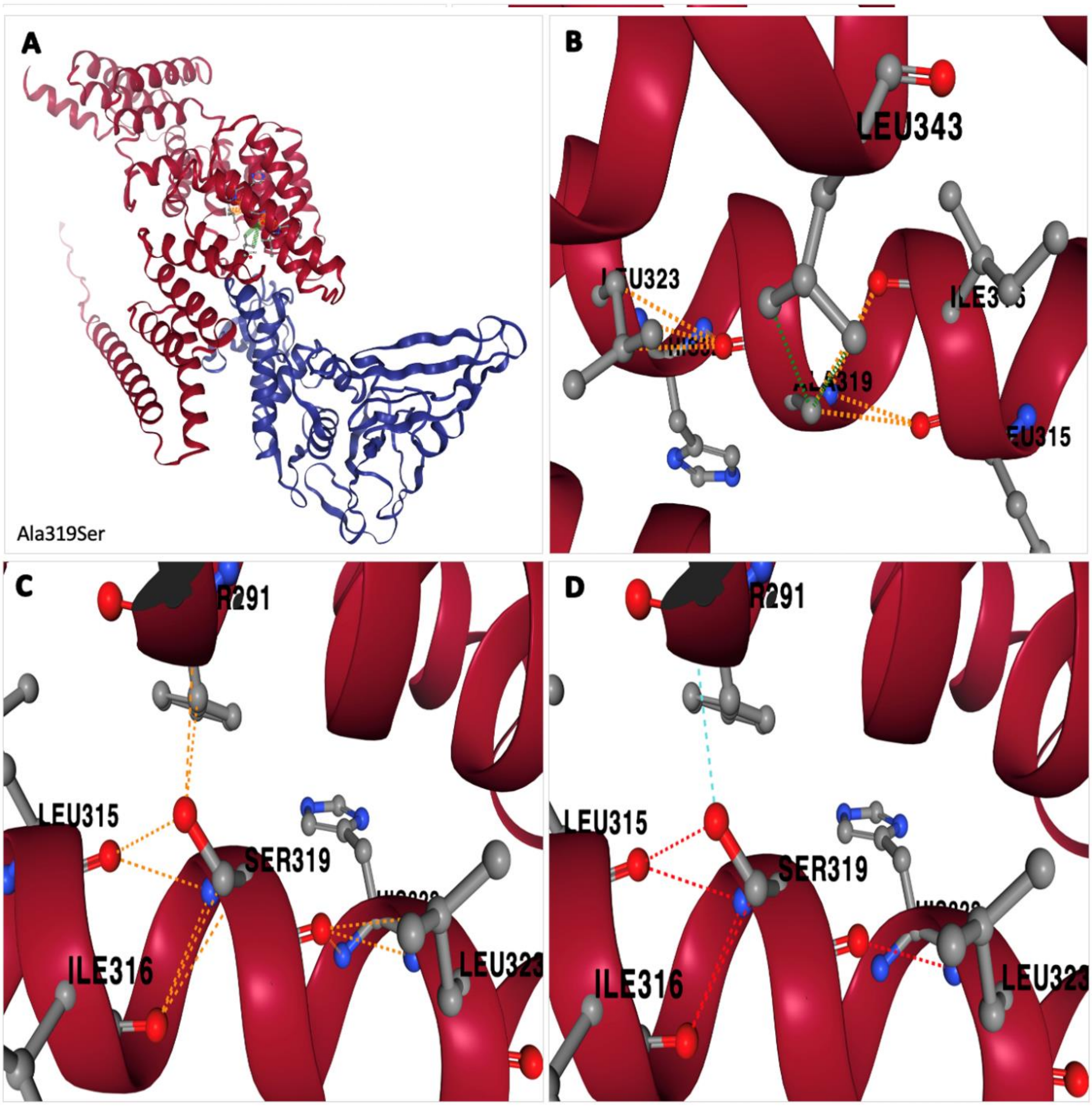

Figure S4_10. Position of residue 319 in IFIT2 and impact of mutation Ala319Ser in IFIT2:SARS-CoV-2 PLpro complex. A) Ribbon model IFIT2:SARS-CoV-2 PLpro complex and the location of residue 319. IFIT2 and SARS-CoV-2 PLpro proteins are shown in red and blue, respectively. B) Illustration of polar and hydrophobic bonds between Ala319 and the neighbouring residues. C, D) Impact of Ala319Ser on the vicinity residues and formation of new hydrogen bonds and Van der Waals contacts. Hydrophobic (green dash lines), Van der Waals (cyan dash lines), polar (orange dash lines) and, hydrogen (red dash lines).

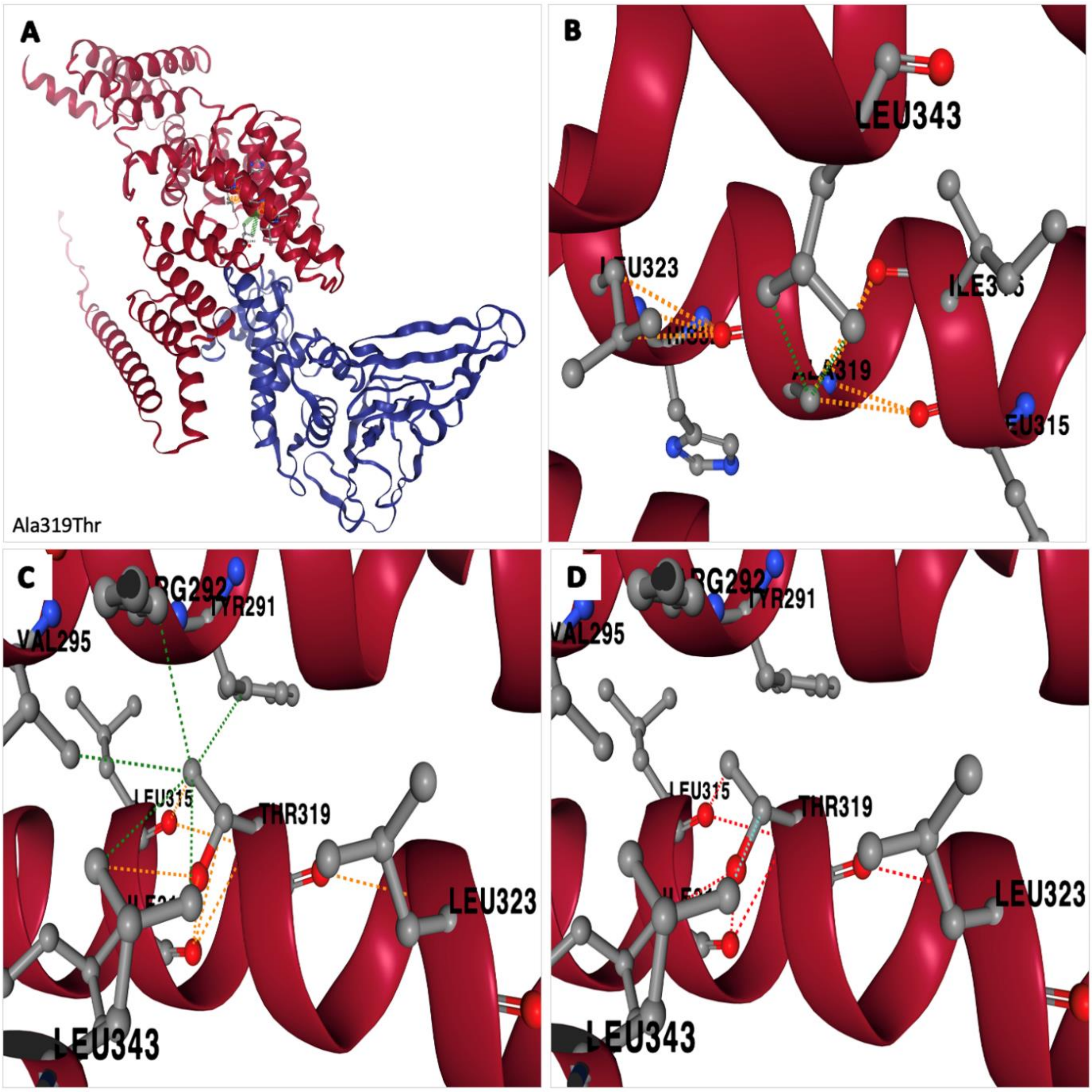


Figure S4_11. Position of residue 319 in IFIT2 and impact of mutation Ala319Thr in IFIT2:SARS-CoV-2 PLpro complex. A) Ribbon model IFIT2:SARS-CoV-2 PLpro complex and the location of residue 319. IFIT2 and SARS-CoV-2 PLpro proteins are shown in red and blue, respectively. B) Illustration of polar and hydrophobic bonds between Ala319 and the neighbouring residues. C, D) Impact of Ala319Thr on the vicinity residues and formation of new hydrogen bonds and Van der Waals contacts. Hydrophobic (green dash lines), Van der Waals (cyan dash lines), polar (orange dash lines) and, hydrogen (red dash lines).

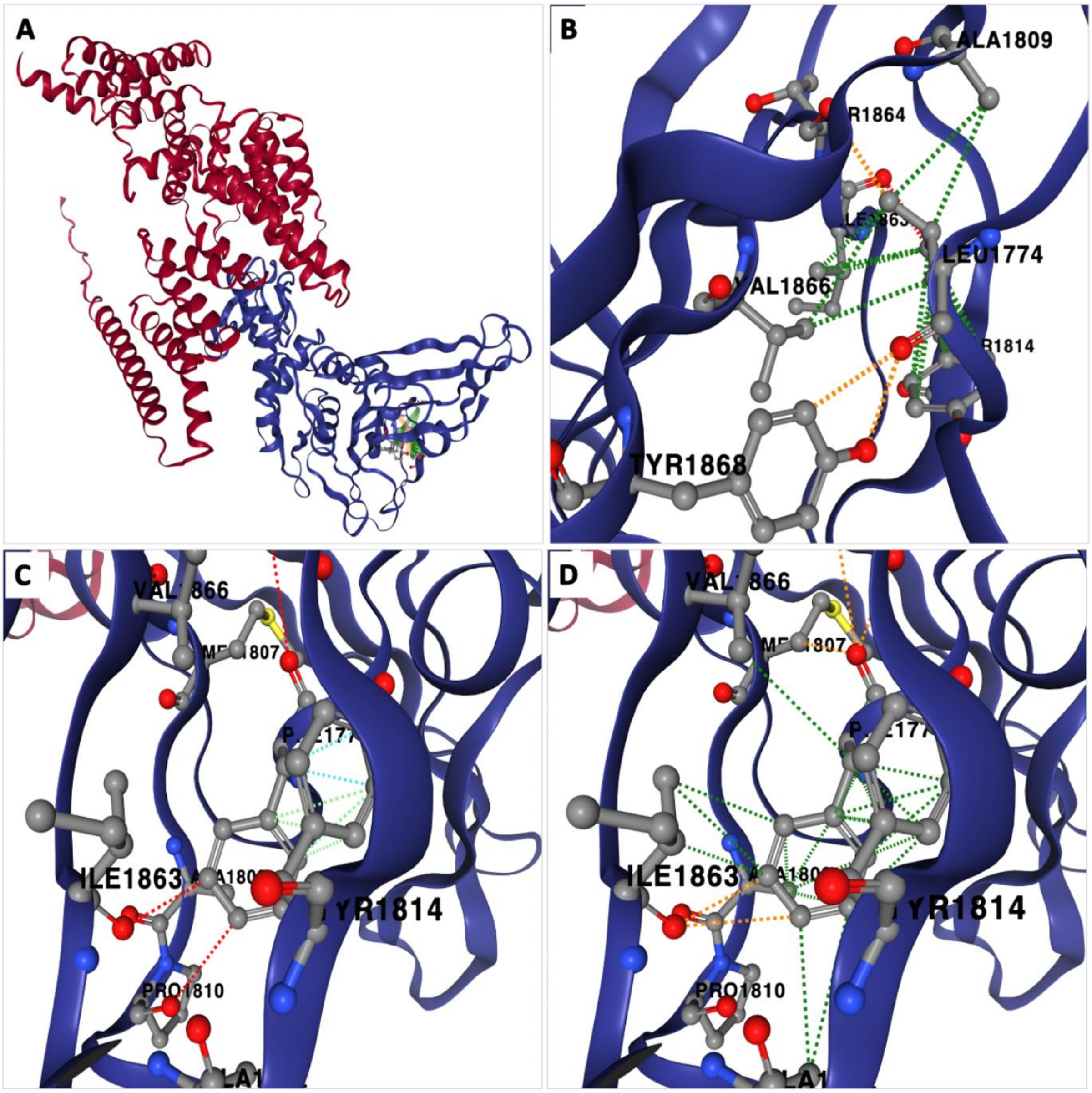


Figure S4_12. Position of residue 1774 in SARS-CoV-2 PLpro and impact of mutation Leu1774Phe in IFIT2:SARS-CoV-2 PLpro complex. A) Ribbon model IFIT2:SARS-CoV-2 PLpro complex and the location of residue 1774. IFIT2 and SARS-CoV-2 PLpro proteins are shown in red and blue, respectively. B) Illustration of polar and hydrophobic bonds between Leu1774 and the neighbouring residues. C, D) Impact of Leu1774Phe on the vicinity residues and formation of new hydrogen bonds and Van der Waals contacts. Hydrophobic (green dash lines), Van der Waals (cyan dash lines), polar (orange dash lines) and, hydrogen (red dash lines).

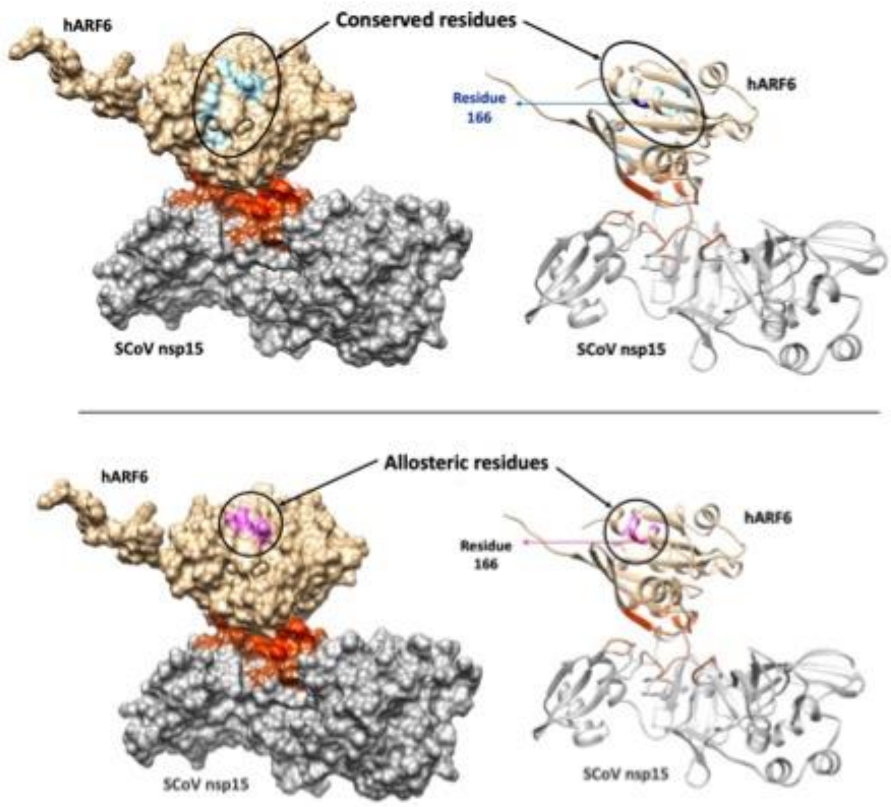


Figure S4_6. Position of hARF6 affinity-enhancing residues: 166. Top) The space-filled (left) and Ribbon (right) models of hARF6:SCoV2 NSP15 complex the position 166 in dark blue and its neighbouring conserved positions in light blue. Bottom) The space-filled (left) and Ribbon (right) models of hARF6:SCoV2 NSP15 complex the position 166 in dark pink and its neighbouring allosteric sites in light purple. Positions 164, 166, 168 and 169 are both conserved and predicted allosteric sites. The red and brown residues are the direct contact residues in hARF6 (tan) and SCoV-2 nsp15 (grey), respectively.
