## Supplementary material for "Predicting human and viral protein variants affecting COVID-19 susceptibility and repurposing therapeutics": Supplementary file 7-Cavityplus-results.docx

1. **CavityPlus results for IFIH1 (green):PLpro (orange) complex**

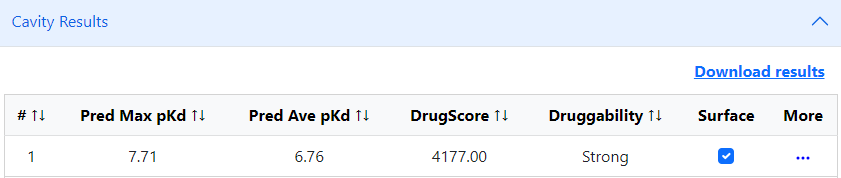

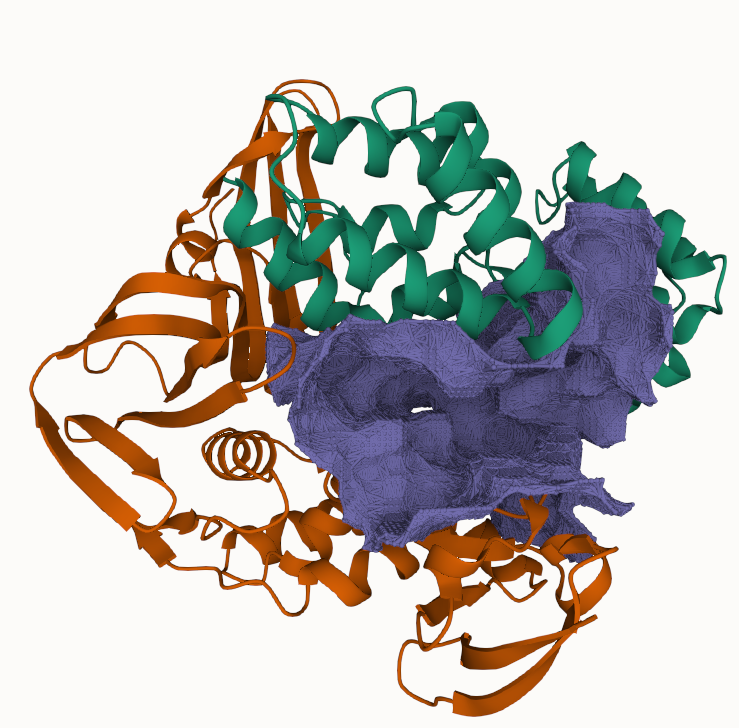

2. Cavityplus results for ARF6 (green):NSP15 (orange) complex

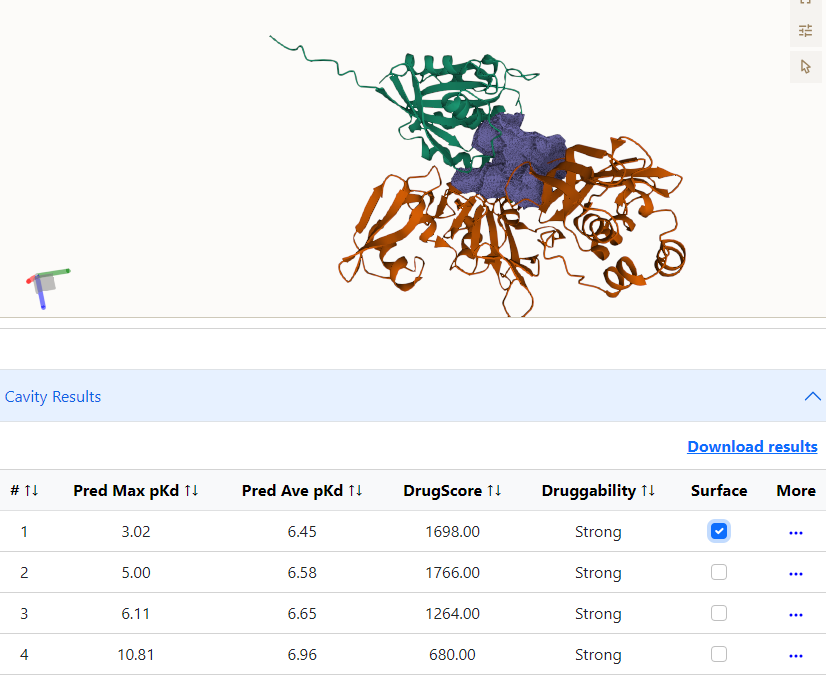

3. CavityPlus results for AXL:NTD complex

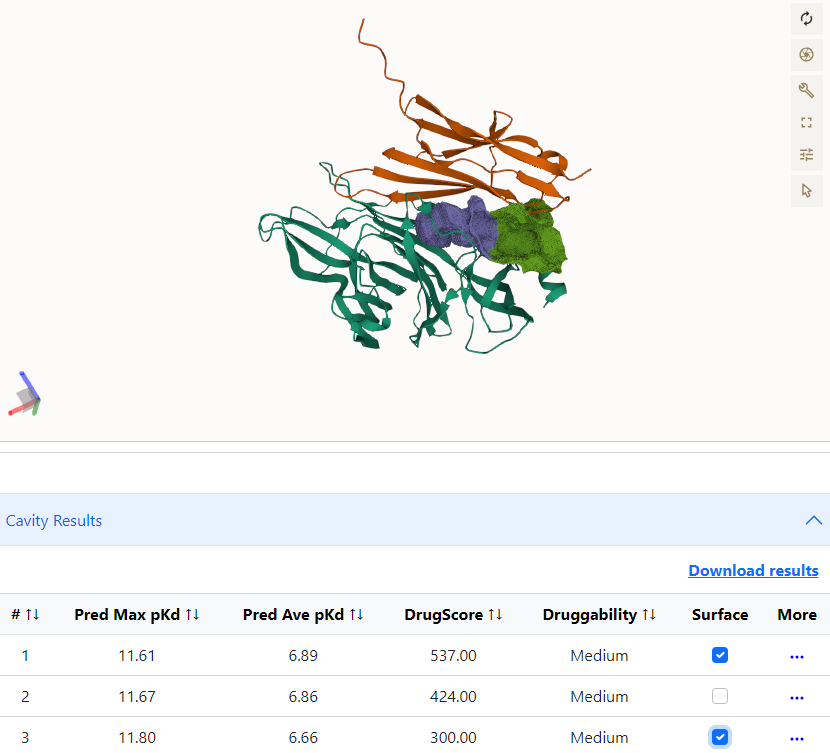

**Source of Cavity figures**: CavityPlus output (<http://www.pkumdl.cn:8000/cavityplus/index.php#/> )

**CavityPlus reference:**

Wang, S., et al., *CavityPlus 2022 Update: An Integrated Platform for Comprehensive Protein Cavity Detection and Property Analyses with User-friendly Tools and Cavity Databases.* J Mol Biol, 2023. **435**(14): p. 168141.
