## Supplementary material for "Predicting human and viral protein variants affecting COVID-19 susceptibility and repurposing therapeutics": Supplementary file 8-dockingIFIH1-plro.docx

**Docking of IFIH1:PLpro complex with ligand with Selgantolimod** (ChEMBL ID: CHEMBL4594258)

**Figure: Docking of** **Selgantolimod with IFIH1:PLpro complex. The ligand binds at the cavity formed by IFIH1:PLpro interface.**

A. Docking of Selgantolimod (blue) with IFIH1 (green) :PLpro (orange) is performed using AutoDock and Autodock Vina. (AutoDock Vina score: -7.4).

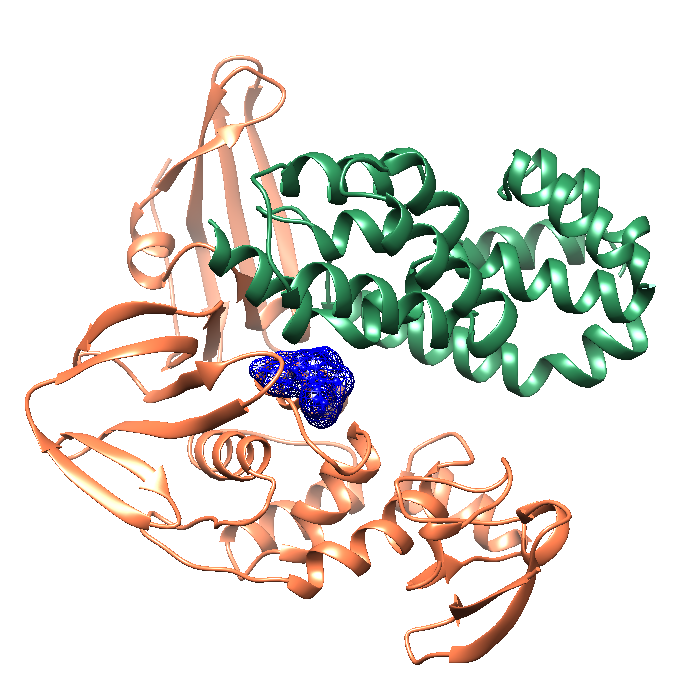

**(B) Selgantolimod binds at the cavity strongly predicted by Cavityplus (score: 4177.0). The cavity residues are shown is purple.**

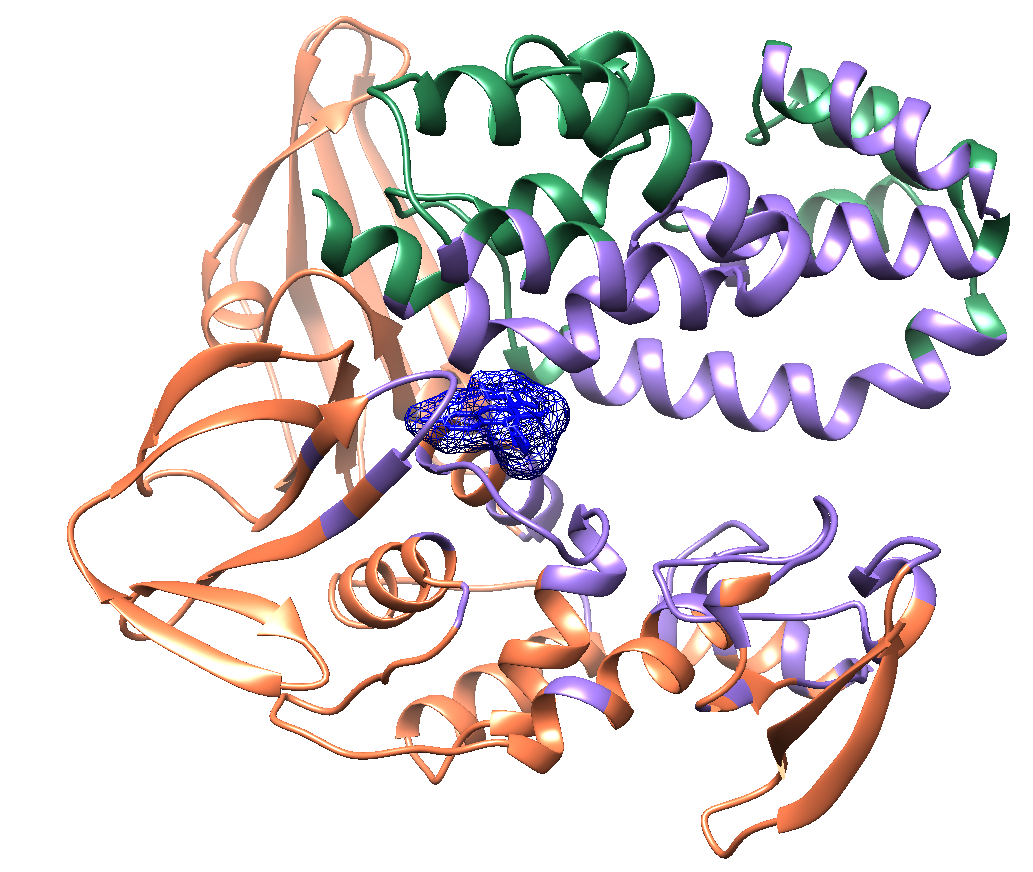

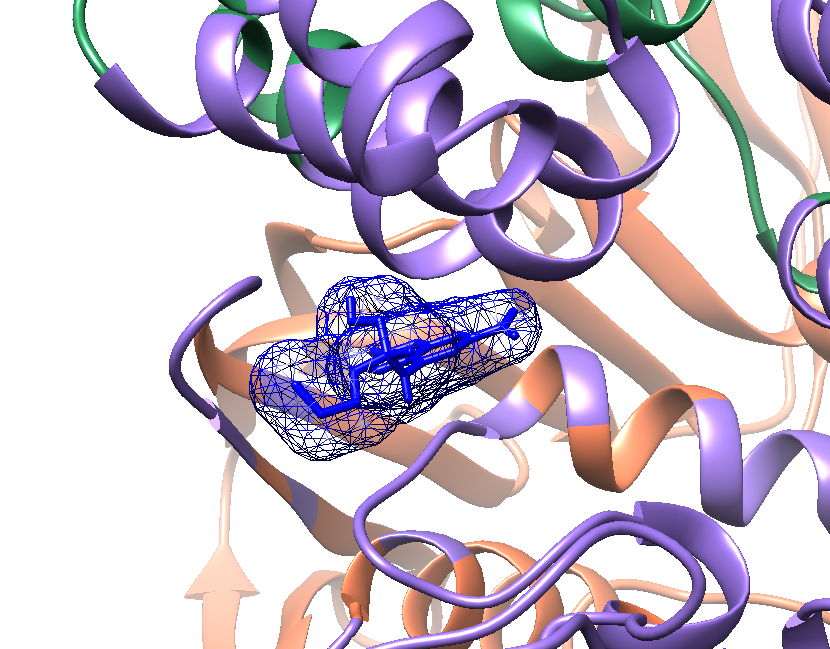
